# Controlled protein activities with viral proteases, antiviral peptides, and antiviral drugs

**DOI:** 10.1101/2023.02.27.530290

**Authors:** Elliot P. Tague, Jeffrey B. McMahan, Nathan Tague, Mary J. Dunlop, John T. Ngo

## Abstract

Chemical control of protein activity is a powerful tool for scientific study, synthetic biology, and cell therapy; however, for broad use, effective chemical inducer systems must minimally crosstalk with endogenous processes and exhibit desirable drug delivery properties. Accordingly, the drug-controllable proteolytic activity of hepatitis C *cis-*protease NS3 and its associated antiviral drugs have been used to regulate protein activity and gene modulation. These tools advantageously exploit non-eukaryotic/prokaryotic proteins and clinically approved inhibitors. Here we expand the toolkit by utilizing catalytically inactive NS3 protease as a high affinity binder to genetically encoded, antiviral peptides. Through our approach, we create NS3-peptide complexes that can be displaced by FDA-approved drugs to modulate transcription, cell signaling, split-protein complementation. With our developed system, we discover a new mechanism to allosterically regulate Cre recombinase. Allosteric Cre regulation with NS3 ligands enables orthogonal recombination tools in eukaryotic cells and functions in divergent organisms to control prokaryotic recombinase activity.

## Introduction

The ability to tune and dynamically control protein activity empowers the study of biological phenomena, engineering of synthetic biology, and design of safer cellular therapies. To achieve protein activity control, small molecules are particularly useful because they can be dose-dependent, dynamic, and can be delivered through multiple administration routes. Prominent examples of such ligand-dependent methods to control protein activity include chemically induced protein proximity (CIPP)^1–5^, induced protein trafficking^6^, and controlled enzymatic activity^7^. However, small molecules used in this context often exhibit endogenous crosstalk in mammalian organisms and the human microbiome, or they exhibit poor bioavailability.

For synthetic biology and future therapeutic applications, an ideal ligand inducible system would be orthogonal to eukaryotic and prokaryotic biology, present at low levels in the environment, versatile, tunable, and dynamic. To employ more orthogonal inducers and build upon the library of existing chemogenetic proteins, we and others developed the ligand inducible connection (LInC) system and stabilizable polypeptide linkages (StaPLs), which utilize Hepatitis C virus (HCV) *cis*-protease NS3 and its host of clinically tested antiviral drugs^8,9^. These systems facilitate control of transcription, protein localization, and cell signaling by utilizing highly specific and characterized ligands developed to bind a non-endogenous target protein.

Yet, an expanded repertoire of ligands exists to inhibit NS3, each with potential value as a synthetic biology tool. Due to the essential function of NS3 in viral replication, multiple approaches have been employed to develop NS3 inhibitor ligands, including genetically encodable ligands such as RNA aptamers and peptides^10,11^. Recent applications utilize NS3 and genetically encoded antiviral peptides to control various protein functions^12–15^. In this schema, NS3 protease serves as a high affinity binder to antiviral peptides, which can be conditionally displaced by small molecule drugs.

Since NS3 in this context serves as a high affinity binder instead of a severable linker, the protease no longer needs to be catalytically active. Although NS3 protease exhibits stringent substrate specificity^16^, there are documented endogenous cleavage sites involved in viral immune response^17,18^, and some recently discovered promiscuity in cleavage sequence^19^. Thus, the ablation of NS3 proteolytic capacity can now be applied to improve the orthogonality of chemosensory tools used for control of protein function.

In this work, we chemically control protein functions using catalytically inactivated or drug-resistant NS3 mutants paired with displaceable, genetically encoded peptides. Through this approach, we expand the utility of antiviral drugs to conditionally regulate transcription, cell signaling, split-protein complementation, and intramolecular inhibition. With our approach, we discover a novel single-chain mechanism to control Cre recombinase with applicability across eukaryotic and prokaryotic domains. Lastly, we discover that specific drug-resistance mutations employed in the proteolytic NS3 tools^9^ extend to peptide displacement tools. We use these properties to create combinatorial drug control of Cre recombinase.

## Results

### Controlled transcription and cell signaling

To first validate our approach, we devised a system to control the assembly of heterodimeric transcription factors with antiviral drugs. We began by fusing a previously developed high affinity antiviral peptide (CP5-46A-4D5E, referred to here as P_MED_)^11^ to the Gal4 DNA-binding domain (Gal4_DB_), and then fused a separate catalytically “dead” NS3^S139A^ serine protease (dNS3) to a transcriptional effector domain (**Fig. 1a**). In the absence of drug, the antiviral peptide and dNS3 are expected to spontaneously assemble to form a functional synthetic transcription factor. Upon the addition of a small molecule drug, dNS3 is expected to competitively bind to drug over peptide, therefore inhibiting activity of the transcriptional effector.

**Figure 1.**
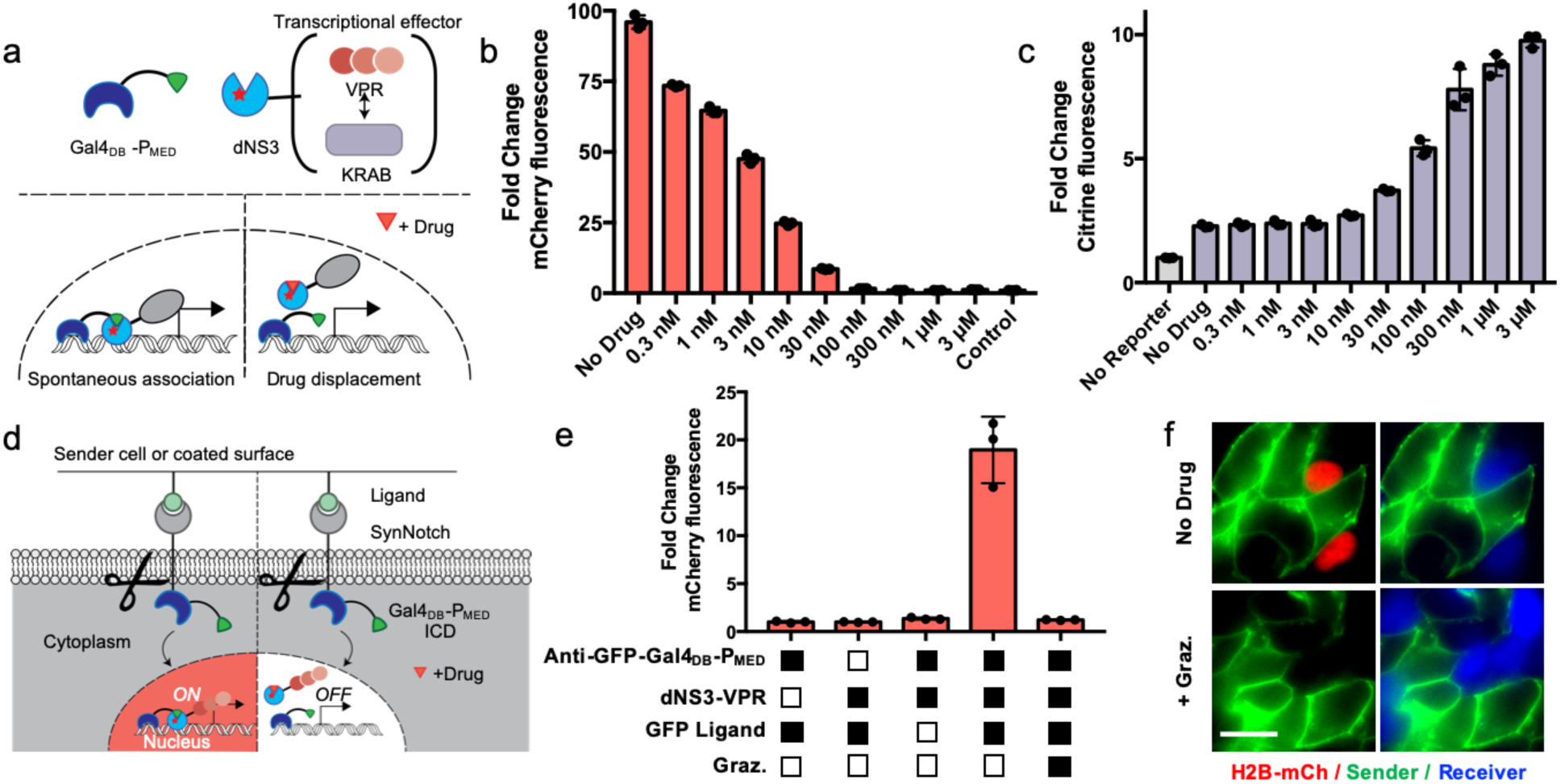
Antiviral drug control of transcriptional effectors and cell signaling. **(a)** Gal4_DB_-P_MED_ complexes with dNS3-fused transcriptional effectors, creating heterodimeric transcription factors that can be competitively inhibited by antiviral drugs. **(b)** Transcriptional turn-off of Gal4 driven H2B-mCherry expression in HEK293FT cells via competitive grazoprevir inhibition of dNS3-VPR binding to Gal4_DB_-P_MED_. **(c)** Drug control of UAS-CMV-H2B-Citrine expression via competitive grazoprevir inhibition of repressor domain dNS3-KRAB binding to Gal4_DB_-P_MED_. **(d)** Design of drug-controlled Notch ICD function. Gal4_DB_-P_MED_ drives expression in the presence of receptor ligand, dNS3-VPR, and absence of drug. **(e)** Combinatorial control of UAS-H2B-mCherry expression via surface coated ligand GFP and drug-controlled displacement of dNS3-VPR from Gal4_DB_-P_MED_ ICD. **(f)** Drug controlled transcription via ligand mediated release of SynNotch Gal4_DB_-P_MED_ ICD in dual HEK293FT coculture. Sender cells are expressing surface presented GFP ligand, and receiver cell contains transiently expressed SynNotch Gal4_DB_-P_MED_ and dNS3-VPR to drive UAS-H2B-mCherry reporter. Receiver cells express iRFP670 (false colored blue) for labeling purposes. Scale bar: 20 μm. All quantitative fluorescent data was captured via flow cytometry of transiently transfected HEK293FT cells, 48 hours post-transfection/drug addition aside from UAS-H2B-mCherry, which was clonally incorporated into HEK293FT. Plotted values are the mean ± SD of biologically independent replicates, n=3. Unless otherwise stated drug refers to 3 μM grazoprevir.

To first mediate transcriptional activity, transcriptional activator VPR^20^ was fused to dNS3. In this configuration, we expected spontaneous association between Gal4_DB_-peptide and dNS3-VPR to lead to transcriptional activation, and drug addition to lead to transcriptional shut-off. Correspondingly, we transiently transfected Gal4_DB_-peptide and dNS3-VPR and observed spontaneous transcriptional activation of a genomically integrated, UAS-H2B-mCherry reporter in HEK293 cells (**Fig. 1b**). As we increased the concentration of the NS3 inhibitor grazoprevir, we observed dose dependent transcriptional inhibition. We tested two peptides developed against NS3^WT^ by Kügler et al. - medium affinity CP5-46A-4D5E, and high affinity CP5-46-4D5E, referred to here as P_MED_ and P_HI_, respectively (**Supp. Table 1**). Interestingly, P_HI_ fused to Gal4_DB_ acted as a transcriptional activation domain even in the absence of dNS3-VPR co-transfection, while Gal4_DB_-P_MED_ displayed no observable activation on its own (**Supp. Fig. 1**). Consequently, only P_MED_ was used for the transcriptional effector work while P_HI_ was reserved for future, non-transcriptional applications. We next predicted that the effector domain could be exchanged to control contrasting transcriptional outputs. To test this, we fused dNS3 to transcriptional repressing domain KRAB to control transcriptional repression of a constitutive reporter gene, UAS-CMV-H2B-Citrine (**Fig. 1c**). As expected, transcriptional repression occurred in the absence of drug, and the extent of repression could be controlled by altering the concentration of grazoprevir.

These transcriptional effectors can also govern programmable cell signaling. To demonstrate this, we sought to apply drug-dependent transcriptional control in tandem with SynNotch cell signaling. SynNotch represents a highly modular approach to sense an extracellular ligand and proteolytically trigger the release of an intracellular domain (ICD). In practice, this customizable cell signaling platform has been used to create multicellular patterning as well as combinatorial antigen-sensing circuits to combat cancer^21,22^. Drug control of SynNotch would enable creation of more complex gene circuits and safety switch behavior for therapeutic applications. For these reasons, we replaced the ICD of a previously developed Anti-GFP SynNotch^21^ with our Gal4_DB_-P_MED_ (**Fig. 1d**). In this arrangement, gene activation is controlled by extracellular ligand induced proteolysis and drug-controlled docking of dNS3-VPR, with the goal of allowing for tight combinatorial control of ICD transcriptional activation of SynNotch.

Cells containing the modified SynNotch constructs were grown on surfaces coated with GFP ligand in the presence of varying grazoprevir concentrations. When presented ligand in the absence of drug, HEK293 cells exhibited UAS-H2B-mCherry reporter activation, while increasing concentration of drug led to dose-dependent transcriptional inhibition (**Fig. 1e, Supp. Fig. 2a**). Due to an extracellular c-Myc epitope tag, transcription could also be controlled by plated c-Myc antibody ligand (**Supp. Fig. 2b**). Small molecule control of SynNotch additionally extended to the application of intercellular signaling, which we tested by creating a GFP surface ligand “sender” cell line and then cocultured these cells with transiently transfected dNS3/peptide SynNotch “receiver” cells. Cocultured receiver cells exhibited drug-dependent transcriptional turn-off when presented GFP ligand (**Fig. 1f, Supp. Fig. 2c**).

**Figure 2.**
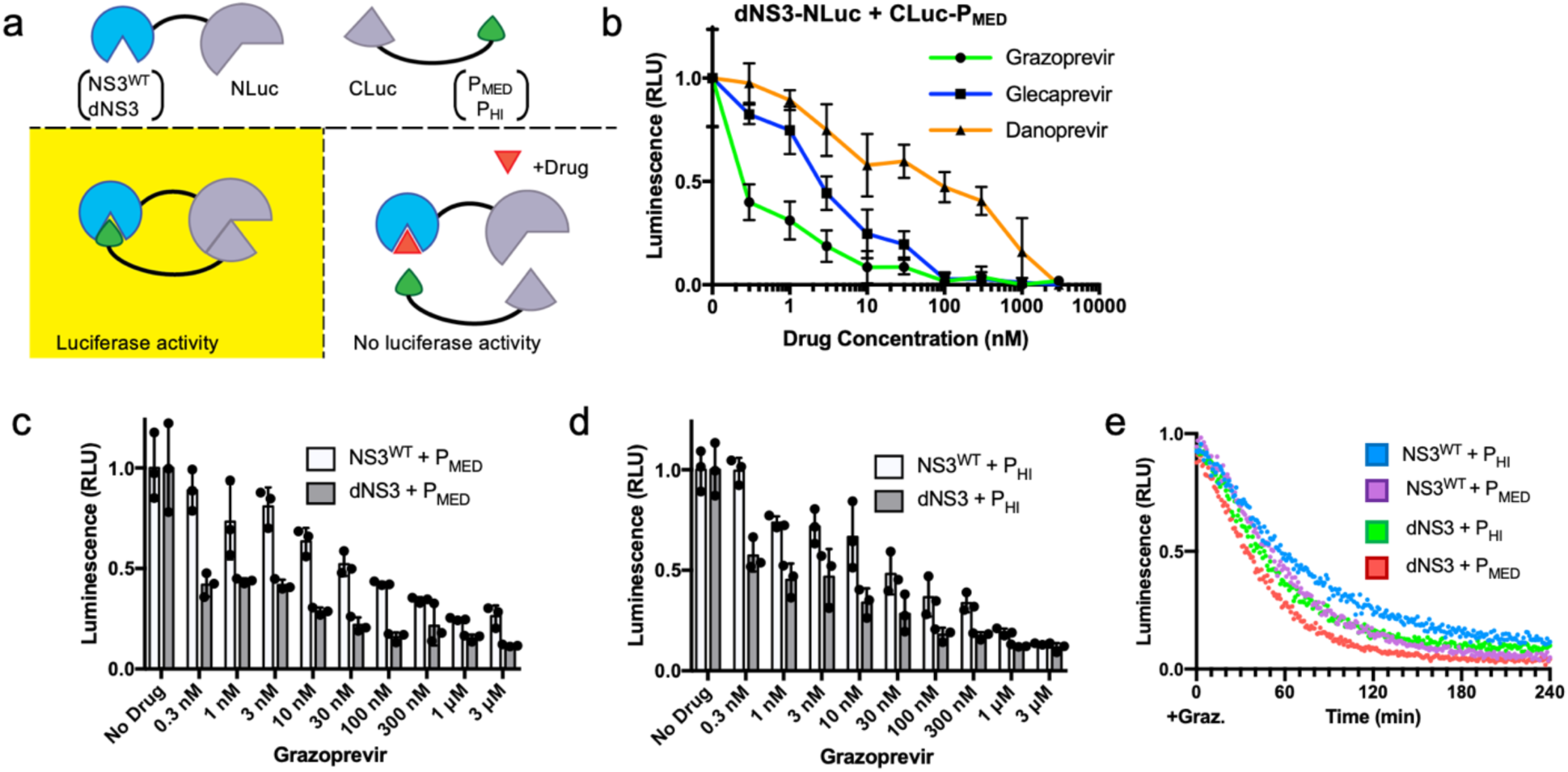
Split Luciferase control and drug dissociation characterization. **(a)** Design of split luciferase constructs. NLuc(12-1247) was fused to dNS3 or NS3^WT^. CLuc(1200-1643) was fused to P_MED_ or P_HI_ peptide. **(b)** Dose response of HEK293FT cells transfected with dNS3-NLuc and CLuc-P_MED_ to various drugs. Lines are connected to guide the eye. **(c)** Dose response of NLuc fused to NS3^WT^ or dNS3 in combination with CLuc fused to P_MED_ or **(d)** higher affinity P_HI_ **(e)**. Temporal response to drug of the four different combinations of NLuc and CLuc fusions to NS3 and peptides. All dose responses are normalized to their respective no drug controls and transfected 48 hours before lysing for luciferase experiments. Drug was added at the time of transfection for dose response experiments. For temporal experiments, 3 μM of grazoprevir was added at the time of lysis. Plotted values are the mean ± SD of biologically independent replicates, n=3. For linear regression and SD of fig 2e, see supplemental info.

### Controlled enzymatic activity by split protein complementation

We next examined the binding characteristics of drug-controlled complementation with the goal of directly demonstrating drug-mediated dissociation of formed NS3-peptide complexes. Since dNS3 and its high affinity peptides effectively regulated assembly of a split transcription factor, we surmised that dNS3/peptide complementation could control assembly of a split protein to control enzymatic activity. To test this, we fused permutations of dNS3, NS3^WT^ and each peptide variant to split fragments of *Renilla* luciferase (**Fig. 2a**). Controlled assembly of split luciferase not only demonstrates enzymatic control, but it can also be used to elucidate binding characteristics through dynamic luciferase complementation assays^23^.

We first turned to the previously identified set of dNS3 and P_MED_ as our model peptide docking pair and tested luciferase activity in transfected HEK293 lysates upon addition of four different drugs: grazoprevir, glecaprevir, danoprevir, and telaprevir (**Fig. 2b, Supp. Fig. 3**). The first three of these drugs inhibit luciferase activity in a dose dependent manner, representing multiple ligand choices with distinct binding profiles. However, split luciferase constructs containing dNS3 exhibit significant resistance to telaprevir. This observation is consistent with the drug’s covalent binding mechanism to the catalytic Ser139^24,25^ which is mutated in dNS3, and this effect is reverted upon replacement with wild-type NS3 (**Supp. Fig. 3**).

**Figure 3.**
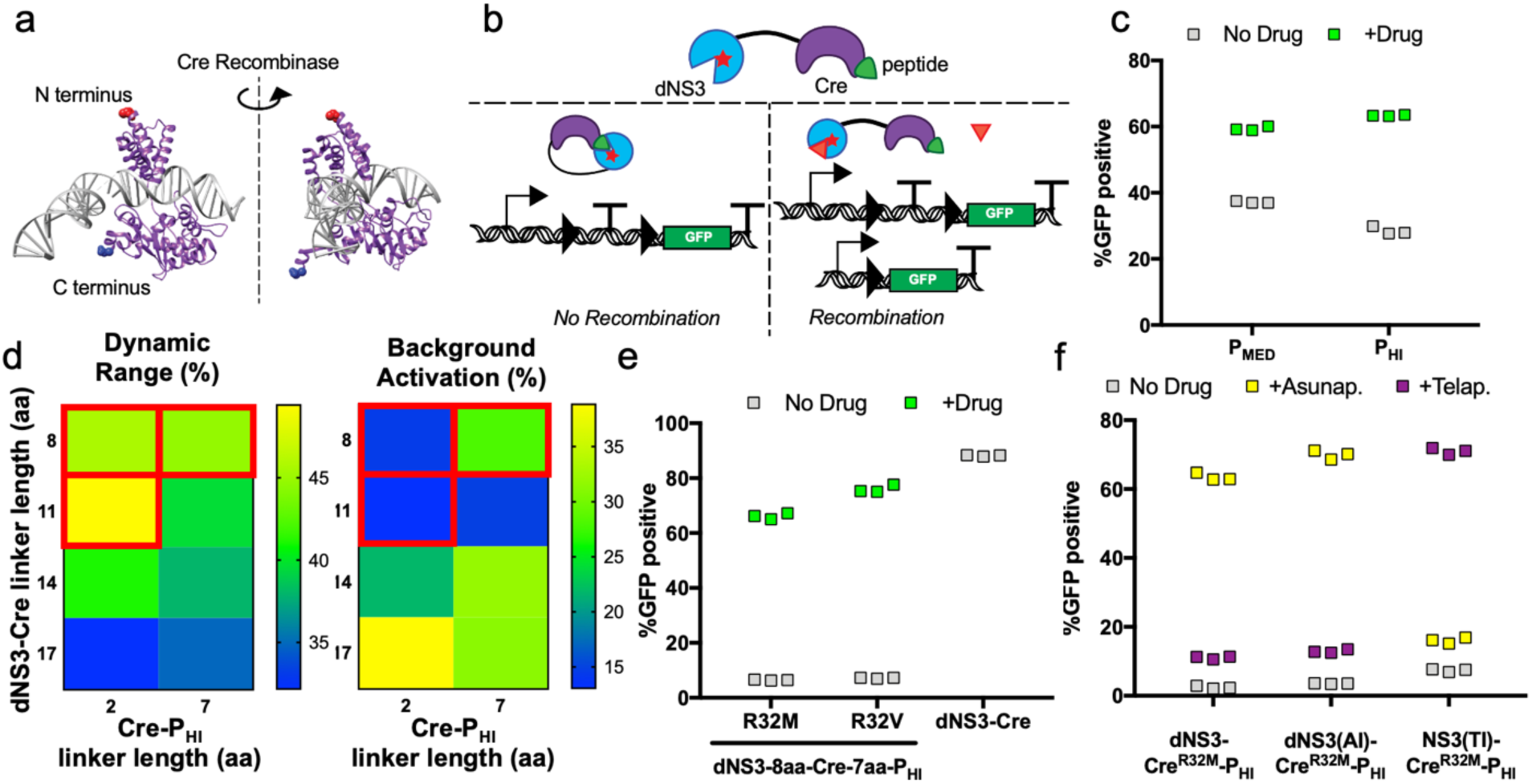
Antiviral drug control of genetic recombination in eukaryotes. **(a)** Structure of Cre recombinase bound to DNA (PDB 1ma7^28^). N-terminal and C-terminal residues labeled in red and blue respectively. **(b)** Design of intramolecular inhibited Cre driving recombination of virally integrated Cre reporter plasmid. Successful recombination results in GFP expression after excision of a stop codon. **(c)** Comparison of transient transfection of dNS3-Cre-peptide with P_MED_ and P_HI_ with flow cytometry, 48 hours post-transfection and drug addition in HEK293FT cells. **(d)** Variation of amino acid (aa) linkers between dNS3-Cre, and Cre-P_HI_ with flow cytometry, 48 hours post-transfection and drug addition. Constructs chosen for highest dynamic range of GFP positive cells upon addition of drug and for low background of GFP positive cells in absence of drug. Optimal linker lengths are boxed in red. **(e)** dNS3-Cre-P_HI_ variants with selected Cre mutants compared against dNS3-Cre control. **(f)** Selectivity of drug resistant NS3-Cre-P_HI_ constructs against asunaprevir (1 μM) and telaprevir (10 μM). Drug was added 24 hours post-transfection, and flow cytometry data was collected 48 hours post-drug addition. Unless otherwise specified, drug refer to 1 μM grazoprevir addition. Plotted values are biologically independent replicates, n=3.

Since NS3’s catalytic serine residue could affect binding characteristics, we tested both NS3^WT^ and dNS3 with P_MED_ and P_HI_ peptides as fusions to split luciferase subunits. All permutations of these constructs were functionally sensitive to the range of grazoprevir concentrations we tested. Notably, the split luciferase heteromerized by dNS3 displays slightly increased sensitivity to drug than NS3^WT^ does (**Fig. 2c**,**d**), possibly due to the loss of a hydrogen bond between the peptides and NS3 as a result of the S139A mutation. However, the affinity differences of the two peptides tested did not seem to make a substantial difference in dose response to grazoprevir at the concentrations tested (**Supp. Fig. 4a**,**b**).

**Figure 4.**
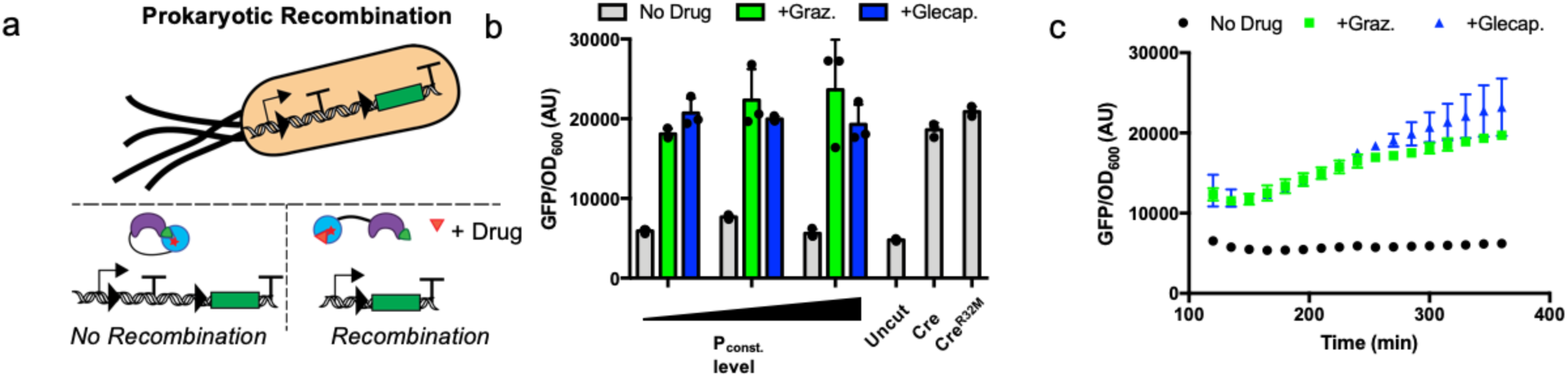
Drug control of genetic recombination in prokaryotes. **(a)** Genetic recombination in prokaryotes. To report on Cre mediated recombination, a transcriptional terminator is located after a constitutive promoter and excised by Cre recombination. **(b)** Constitutively expressed dNS3-Cre^R32M^-P_HI_ in *E. coli* with no drug treatment compared to 50 μM of grazoprevir or glecaprevir treatment, six hours post drug addition, fluorescence/OD quantified by plate reader. Three constitutive promoters (P_const._) were tested with low to medium level expression. These inducible Cre constructs are compared to lowly expressed constitutive Cre and Cre^R32M^. **(c)** Time course response of dNS3-Cre^R32M^-P_HI_ in *E. coli* during exponential growth with and without drug treatment (50 μM, added at time 0). Values are shown starting after 120 min due to low OD prior to this time. Values shown are fluorescence (a.u., superfolder GFP) normalized to cell density (OD_600_), mean ±SD, n=3 biologically independent replicates.

We next demonstrated the temporal displacement of peptide by taking advantage of the reversibility of split luciferase complementation. Temporal dissociation of proteins is important for many biological processes and is often desired for dynamic protein activity control. Kinetics of displacement were measured over a four-hour period with excess drug concentration and normalized to their respective no drug controls (**Fig. 2e, Supp. Fig. 5a**,**b**). In this configuration, the fusions of dNS3 and P_MED_ to split luciferase yielded the fastest displacement kinetics, with luciferase activity reduced to half maximal activity at ~30 minutes. The replacement of dNS3 with NS3^WT^ or the replacement of P_MED_ with P_HI_ increased the time to displacement, reflecting expected increases in protein-peptide affinity. Following peptide and NS3 variations, we tested multiple drugs with dNS3 and P_MED_ (**Supp. Fig. 6a**,**b**), but observed little difference in dissociation rate at saturating drug concentrations aside from telaprevir, which did not dissociate dNS3-P_MED_ complexes well. Altogether, the classes of previously developed peptides and small molecules allow for variation of binding kinetics and dose responses with multiple antiviral drugs. Temporal experiments additionally reveal that antiviral drugs can disrupt preformed protein-peptide complexes.

### Controlled genetic recombination through intramolecular inhibition

Disruption of preformed protein-peptide complexes led us to hypothesize that we could control intramolecular protein interactions, i.e., engineer synthetic protein allostery. In native biology, enzymes are routinely controlled by allosteric mechanisms. Synthetic protein allostery is advantageous in a biological tool because it offers a self-encoded mechanism of control, and it can expand the utility of a protein tool across organisms. To demonstrate these capabilities, we next sought to allosterically control enzyme activity via dNS3 and an inhibitory peptide.

For this application, we focused on Cre recombinase because it is a widely used enzyme that often requires ligand control. Cre recombinase is a site specific, DNA recombinase that has been traditionally controlled by conditional nuclear translocation via 4-hydroxytamoxifen^7^ or various split protein complementations^26^. Yet, Cre is commonly used in developmental biology studies, which often require genetically small constructs for viral incorporation and orthogonal inducer ligands that are lowly present in the environment. For these types of applications, we instead envisioned a single construct, drug inducible Cre controlled by antiviral drugs. Cre is a tyrosine recombinase, and since no available unbound Cre structures existed, we compared the structure of bound Cre (**Fig. 3a**) to related unbound and bound tyrosine recombinases such as XerA and XerD^27,28^. Comparisons reveal that each of these tyrosine recombinases forms a C-shaped clamp composed of two domains, separated by a flexible linker. Furthermore, recent studies of Cre recombinase demonstrate independent folding mechanisms of the two N- and C-terminal domains^29^. Based on these observations, we inferred that Cre undergoes a large conformational change to regulate DNA cleavage and hypothesized that the fusion of dNS3 and inhibitory peptide at opposing termini of Cre could result in drug controlled intramolecular occlusion of the enzyme (**Fig. 3b**). To measure drug inducible dNS3-Cre-peptide recombination, all constructs were tested in a clonal Cre-stoplight reporter HEK293 cell line in which Cre activity results in constitutive GFP expression. Initial testing with P_MED_ peptide revealed promising results, and dynamic range improved with replacement of P_MED_ with higher affinity P_HI_ (**Fig. 3c**).

Recombination modulates expression of GFP with a binary DNA excision event, making it crucial to reduce basal recombination in the absence of drug. Following substitution of P_MED_ with P_HI_, we began further optimization for lower background recombination while retaining high dynamic range. To further reduce basal recombination, flexible amino acid (aa) linker length was varied between the N-terminal fusion of dNS3 to Cre and between the C-terminal fusion of P_HI_ to Cre (**Fig. 3d**). From the linker variations tested, we chose three candidates with high dynamic range of activation upon grazoprevir addition and low background activation in absence of drug (**Fig. 3d, boxed in red**).

Before moving forward with these three selected linker variants, we noted that there was still significant background activation, which we aimed to reduce with further optimization. We hypothesized that some intramolecular dNS3 and P_HI_ interactions were spontaneously active, subsequently allowing these Cre molecules to cooperatively dimerize through native Cre termini interactions^30,31^. To mitigate cooperativity and decrease background, we next mutated Cre R32 site, which is thought to decrease cooperativity and has been shown to increase recombination accuracy^32^. This mutation choice is further supported by recent development to control Cre with light inducible AsLOV2 and destabilizing N- and C-terminal Cre mutations (coined LiCre)^33^. The Cre^R32M^ mutation caused considerable decrease in background for all dNS3-Cre-P_HI_ constructs tested in transient transfection (**Supp Fig. 7a**), and linker variant dNS3-8aa-Cre^R32M^-7aa-P_HI_ provided optimally low background recombination and high dynamic range. This linker variant exhibited favorable properties with either Cre^R32M^ or similarly reported Cre^R32V^ mutations, suggesting that this mutation site results in a drug inducible Cre with minimal background in addition to its previously characterized higher accuracy of recombination (**Fig. 3e, Supp. Fig. 7b-d**). Each of these final inducible Cre constructs activated a similar fraction of cells compared to their Cre^WT^ counterparts, while the dNS3-Cre^WT^ control remained constitutively active (**Supp. Fig. 7c, d**).

Equipped with an optimized inducible Cre system, we explored opportunities to control of genetic recombination in two different cell populations to reflect the higher complexity of gene expression in specific cell lineages. To accomplish this, we sought to make orthogonal inducible Cre recombinases. In previous work, Jacobs et al. demonstrated that specific mutations to active NS3 protease could create drug resistant orthogonal protease pairs that are asunaprevir inducible (AI) or telaprevir inducible (TI)^9^. We hypothesized that the antiviral peptides would retain reasonable affinity to mutated NS3 proteases due to the large surface area peptides bind on NS3, and subsequently these NS3 mutants could be used in our NS3-peptide control systems. Introducing these protease mutants into our peptide-based inducible Cre system led to a new pair of orthogonal Cre constructs that can be strongly induced via asunaprevir or telaprevir with low background recombination (**Fig. 3f and Supp. Fig. 8a, b**). Alternatively, the original catalytic NS3^S139A^ mutation is also sufficient on its own to reduce telaprevir sensitivity while retaining asunaprevir induction (**Fig. 3f**), consistent with previous experimental results (**Supp. Fig 3**). Applying the NS3^S139A^ mutation to the NS3(TI) similarly ablates telaprevir sensitivity, leaving only sensitivity to grazoprevir among the three drugs tested here. Collectively, these inducible Cre recombinases can respond to two distinct antiviral drugs and also demonstrate a mechanism of covalent ligand control of enzymatic activity that can be reversed with catalytically inactive NS3.

Lastly, we recognized that drug-controlled recombination with a single-chain Cre construct could potentially translate across divergent organisms. Given our engineered control mechanism of Cre does not rely on nuclear translocation and instead relies on allosteric control (**Supp. Fig 9**), we expected antiviral drug inducible Cre to be highly versatile across organisms and function in prokaryotes. To test this, we co-transformed dNS3-Cre^R32M^-P_HI_ and the GFP Cre reporter into *Escherichia coli* (**Fig. 4a**). Cre constructs were expressed at a constant level under an arabinose inducible promoter (**Supp. Fig. 10a**) or under a broad range of constitutive expression levels (**Fig. 4b, Supp. Table 2**). Upon induction with antiviral drugs, GFP expression significantly increased, with detectable GFP expression occurring at <120 min under a constitutive promoter (**Fig. 4c**) and no significant change to bacterial growth (Supp. Fig. 10b, c). We observed minimal background recombination via dNS3-Cre^R32M^-P_HI_ in the absence of drug, while in contrast, both dNS3-Cre^R32M^ lacking inhibitory peptide and Cre^R32M^ exhibited substantial recombination. The dNS3-Cre^R32M^ and Cre^R32M^ background recombination could not be remedied even under an uninduced, arabinose-driven promoter (**Supp. Fig. 10a**). Interestingly, drug concentrations of antiviral drug inducible Cre needed to be increased from the levels used in mammalian cells, consistent with previous literature utilizing HCV NS3 in bacteria^34,35^. Nevertheless, maximal recombination occurred as low as 6.25 μM of glecaprevir or grazoprevir (**Supp. Fig. 11**). Overall, this tool enables drug inducible recombination in prokaryotic cells and mammalian cells with chemical inducers that are lowly toxic to both cell types.

## Discussion

In this study, we demonstrate diverse controls over protein activity via dNS3, genetically encoded inhibitory peptides, and highly characterized antiviral drugs. To this end, we show antiviral drug regulation of transcriptional complex formation, cell signaling, split enzyme domains, and synthetic allostery of Cre recombinase.

Analysis of peptide binding characteristics revealed tunable and dynamic control of protein-protein interaction, and this system can respond to a myriad of different drugs. Dose responses of these systems can also be tuned via swapping genetic parts such as peptides, alternate forms of NS3, or by the specific drug used. These features underscore the benefits to using a highly characterized target such as NS3 as synthetic tool. Due to the diversity of approaches to target NS3, high affinity, bioavailable small molecule drugs can be paired with alternate inhibitors such as genetically encoded peptides. Furthermore, the effort to develop ligands against NS3 is ongoing^36^, with FDA approvals of molecules such as glecaprevir as recently as 2017, providing researchers a continually expanding toolkit to control these protein tools synthetically.

Alongside our demonstration of the versatility of NS3 and its associated inhibitors, we devised a new mechanism to allosterically control Cre recombinase through structure-guided design. This system displayed the ideal characteristics of low background recombination and high dynamic range while utilizing orthogonal ligands and a single optimized protein. Extensive clinical efforts to develop NS3 inhibition enabled us to expand the repertoire of Cre recombinase tools as well. Building upon characterization of drug-resistant mutants and proteolytic tools^9^, we extended drug-resistant NS3 mutants to control peptide binding with two orthogonal, FDA approved inhibitors.

Lastly, allosterically controlled Cre and antiviral drug inducers easily transferred across domains of life to function in *E. coli*. Minimal toxicity to prokaryotic and eukaryotic hosts opens up greater possibilities of studying causal roles of microbiome gene expression *in vivo* as well as engineering industrial bacterial processes. In contrast to many available bacterial chemical inducers, NS3 inhibitors are not known metabolites of bacterial cells, which increases ligand orthogonality for future microbiome applications. As a whole, NS3 and its associated inhibitors represent an exciting toolkit to control protein activity in a more orthogonal manner for eukaryotic and prokaryotic organisms.

## Acknowledgements

Support for this was provided through NIH research grants R35 GM128859 (to J.T.N.) and R01 HL147585 (to J.T.N.). E.P.T. was also supported through a T32 NIH Training Grant awarded to Boston University (EB006359). M.J.D and N.T. acknowledge funding from NIH Grant R01AI102922 and DOE Grant BER DE-SC0019387. The pEV-UAS-H2B-Citrine reporter plasmid were gifts from M. Elowitz (Caltech, Pasadena, CA, USA). The Cre Reporter plasmid was a gift from N. Geijsen (Leiden University, Netherlands). The pCAG-ERT2CreERT2 plasmid was a gift from C. Cepko (Harvard University, MA, USA). We also thank Ngo lab members for suggestions on the manuscript.

## Contributions

E.P.T and J.T.N devised the concept and design. E.P.T, J.B.M. and N.T. performed and analyzed experiments. J.T.N. and M.J.D. supervised the work. E.P.T. and J.T.N. wrote the manuscript and prepared figures with input from J.B.M., N.T. and M.J.D.

## Declaration of Interests

A patent application has been filed based on this work (J.T.N, E.P.T., N.T., and M.J.D.).

## Methods

### DNA Constructs

Standard procedures of ligation and Gibson assembly were applied to all constructs created for this paper. The Cre reporter plasmid (Addgene #62732) was a gift from Niels Geijsen (Hubrecht Institute), the eGFP-pBAD plasmid (Addgene #54762) was a gift from Michael Davidson (Florida State University), and the pmiRFP670-N1plasmid (Addgene #79987) was a gift from Vladislav Verkhusha (Albert Einstein College of Medicine). See supplemental note for annotated DNA sequences used in this work.

### Mammalian cell culture

HEK293FT cells were purchased from Thermo Fisher and maintained in a 37°C incubator supplemented with 5% CO_2_. Cells were cultured in DMEM with 10% FBS and supplemented with non-essential amino acids (Life Technologies) and Glutamax (Life Technologies). For UAS-H2B-mCherry experiments, a clonal cell line was previously created via random plasmid insertion and serial dilution to select single colonies. Cells were selected for with Zeocin resistance (100 μg/mL, Invivogen). For generation of a Cre reporter line, the lentiviral Cre reporter plasmid (62732) was modified to have Blasticidin resistance. This construct was then virally transduced and selected for with Blasticidin (10 μg/mL, Invivogen) and clonally selected by serial dilution.

### DNA transfections and NS3 inhibitors

All DNA transfections were carried out using Lipofectamine 3000 (Thermo Fisher) according to the manufacturer’s instructions. All transcriptional effector experiments were carried out at a mass ratio between 1:2 and 1:3 of Gal4_DB_ constructs to transcriptional effector. For imaging experiments, fibronectin (Product #F1141, Sigma-Aldrich) was seeded on to glass coverslip substrate at a dilution of 5 μg/mL for one hour at room temperature and rinsed three times with PBS before seeding cells. All other experiments were carried out on tissue cultured treated plastic (Corning) or for cell adherence by manufacturer (ibiTreat by Ibidi).

Drug was either added at time of transfection or 24 hours later. Grazoprevir, danoprevir, and glecaprevir were all purchased from MedChemExpress. All stocks were diluted between 3-10 mM stock concentrations in DMSO.

### Protein Purification

GFP SynNotch ligand was expressed and purified from *E. coli* DH10B cells (Thermo Fisher). *E. coli* were transformed with arabinose inducible EGFP-pBAD plasmid (Addgene #54762) overnight at 37°C. The following day, the bacterial culture was diluted 1:50 into 0.5 L of LB and induced at an OD600 of 0.8 with 0.02% arabinose for 24 hours at 25°C. After induction, bacteria were pelleted at 3000 rcf and purified following the QiaExpress Ni-NTA Fast Start protein purification protocol (Qiagen, Catalog # 30600) under native conditions.

### Ligand coating and coculture for SynNotch experiments

For plated ligand experiments, untreated tissue culture plastic was incubated with purified GFP ligand or c-Myc Monoclonal Antibody (Invitrogen 9E10.3, Catalog # AHO0062) in PBS for one hour at room temperature. Plates were subsequently washed three times with PBS prior to before adding transfected cells and drug mixed in cell culture media.

For coculture experiments, sender cells were virally transduced with constitutive surface presenting GFP-ligand and allowed to grow for two days. Lentivirus was prepared via transient transduction of GFP-ligand construct, VSVG and PAX2 packaging plasmids for 6-12 hours before removing transfection media. After 24 hours and 48 hours, media was collected and replaced. The supernatant was then filtered with 0.45 μM sterile filters (Whatman, Catalog # 6780-2504). Suspended cells were then transduced with viral media for 48 hours before proceeding with coculture. iRFP670 labeled receiver cells were transiently transfected with SynNotch constructs 24 hours prior to coculture. Cells were added at ratios between 1:4 and 1:10 receiver cells to sender cells and cultured for 24 hours before fixation in 4% paraformaldehyde.

### Flow cytometry

Cells were analyzed by flow cytometry in suspension on an Attune NxT flow cytometer (Thermo Fisher) and analyzed by FlowJo (v10). Live cells were gated by forward and side scatter detection. Of the live cells, populations were gated for fluorescently positive if their fluorescence intensity was greater than or equal to the top 1% of non-transfected cells. For Gal4-DB-peptide and NS3-transcription effector experiments, a ratio of 1:2 molar ratio of DNA binding domain to transcriptional effector was used. A co-transfection marker of a constitutively expressed protein, mTurq2, was used and cells were deemed positively transfected if they were mTurq2 fluorescently positive. Hereafter, the geometric mean of reporter UAS-H2B-mCherry (dNS3-VPR experiments) or UAS-CMV-H2B-Citrine (dNS3-KRAB experiments) was measured and normalized to a control.

For Cre recombinase control in mammalian cells, live cells were gated in the same manner as above and gated for transfection by either a co-transfection marker of pmiRFP670-N1 or inter-plasmidic constitutive expression of iRFP670 under a separate constitutive promoter. Cells were further deemed as GFP positive if their fluorescence intensity was greater than or equal to the top 1% of non-transfected cells. Baseline activation of GFP positive cells was then subtracted using transfection marker only cells as baseline for all measurements.

### Luciferase assay

HEK293FT cells were transfected with split luciferase constructs. In the case of drug pre-treatment for dose curve experiments, drug was added at the indicated concentration at the time of transfection. Two days later, the transfected cells were lysed with 1x Ex Luciferase Assay Buffer using the Nano-Glo® Dual-Luciferase® Reporter Assay System (Promega). Immediately after, the samples were treated with an additional volume of 1x Ex Luciferase Assay Buffer with the indicated concentration of drug. The samples were then read with a plate reader once per minute measuring luminescence with 500 millisecond exposure time. Samples which had not been incubated in drug had decreased luminescence over time, corresponding to utilization of the luciferase reagents. For kinetics analysis, linear regression analysis was performed on the untreated samples and used to normalize the data for the treated samples.

### Bacterial cell culture

*E. coli* strain BW25113 was used for all bacterial experiments. Drug inducible Cre expression plasmids were derived from the low copy (SC101) plasmid pBbS8c from Lee et al.^37^. For readout of Cre recombination, a transcriptional terminator flanked by LoxP sites was located between a constitutive promoter and the gene encoding superfolder GFP (sfGFP). The terminator is excised upon recombination, allowing transcription of sfGFP. Cells were transformed and inoculated in 3mL LB with 25 µg/mL chloramphenicol and 30 µg/mL kanamycin for Cre expression and reporter plasmid maintenance, respectively. Cells were grown at 37°C with 200 rpm shaking. In the case of arabinose inducible constructs (**Supp. Fig. 10**), cultures were pre-treated with 1mM arabinose for 2 hours prior to drug treatment when appropriate. Experiments were carried out in clear 96-well plates with cell cultures diluted 1:100 into a culture volume of 200 μL. OD_600_ and fluorescence were measured in a BioTek Synergy H1 plate reader after 6 hours of drug treatment. For sfGFP detection, excitation and emission wavelengths of 480nm and 510nm were used, respectively.

### Image acquisition and analysis

To prepare cells for imaging, transfected cells in ibiTreat 8-wells (iBidi) were fixed in 4% paraformaldehyde (Thermo Fisher) diluted in PBS for less than or equal to ten minutes, followed by three rinses with PBS, and then quenching with 5% BSA. Fixed cells were maintained and imaged in PBS.

Images were acquired with epifluorescence using a Zeiss AxioObserver.Z1 microscope and Zen software (Zeiss). Representative images were taken and processed in ImageJ-based image analysis package Fiji.

### Statistics

All statistical analyses for flow cytometry were performed in Prism (v7.04) using three independent biological replicates. For imaging conditions ≥3 images were taken per condition and representative images were selected. Statistical significance was determined via standard t-tests in Excel. For temporal luciferase experiments, linear regressions were also performed in Prism (v7.04) using three independent biological replicates.

