## Supplementary Material for "Controlled protein activities with viral proteases, antiviral peptides, and antiviral drugs"

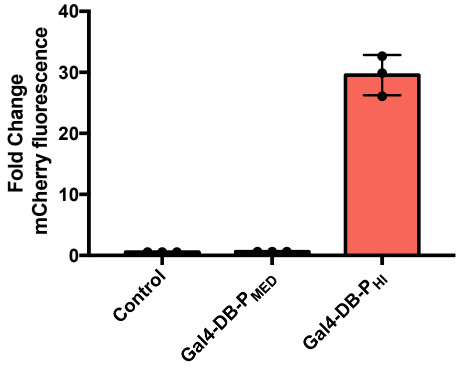


**Supplementary Figure 1 | $Gal4_{DB}$-$P_{HI}$ acts as a transcriptional activator.** Transcriptional activation of UAS-H2B-mCherry via $Gal4_{DB}$-$P_{MED}$ or $Gal4_{DB}$-$P_{HI}$ without drug or dNS3-VPR compared to control of dNS3-VPR only. Plotted values are the mean ± SD of biologically independent replicates, n=3.

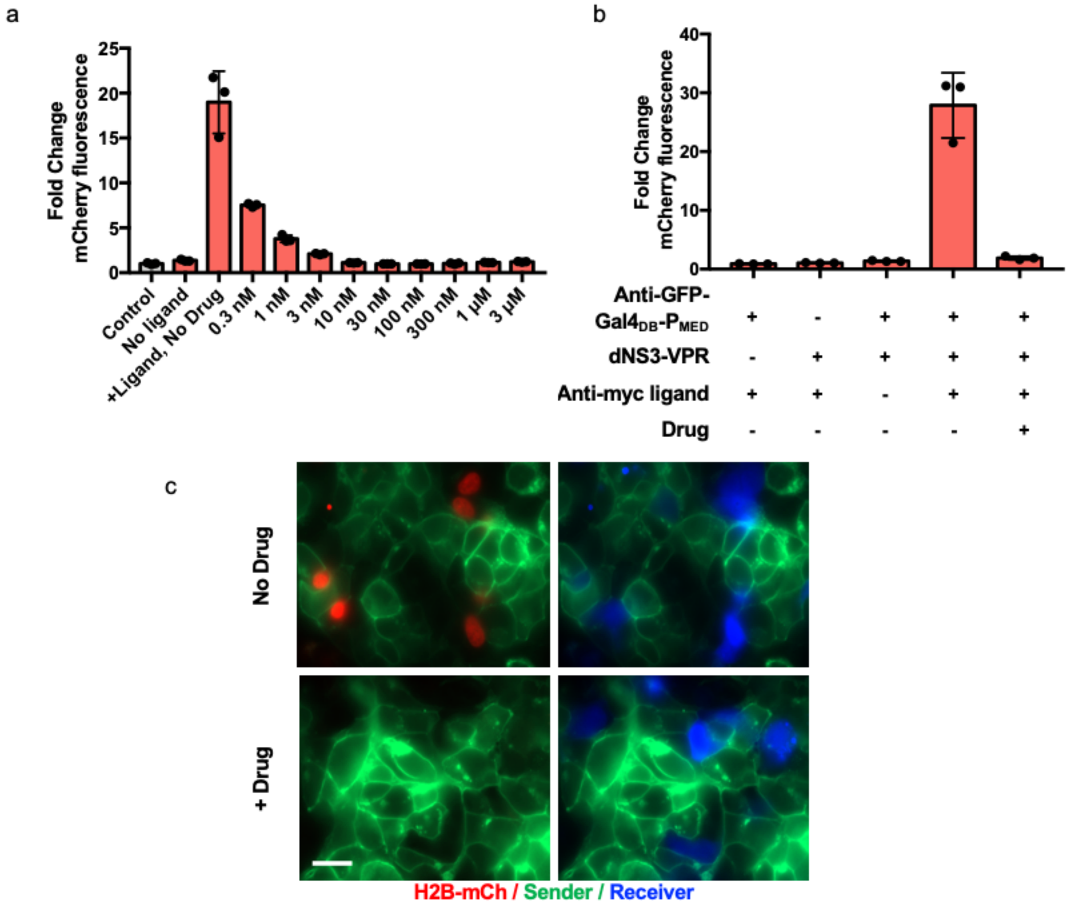


**Supplementary Figure 2 | SynNotch with a Gal4$_{DB}$-P$_{MED}$ ICD displays dose dependent activation and can respond to GFP or anti-myc ligand. (a)** Dose dependent response of transcriptional activation via ligand mediated release of SynNotch Gal4$_{DB}$-P$_{MED}$ ICD and varying amount of NS3 inhibitor, grazoprevir, compared to anti-GFP-Gal4$_{DB}$-P$_{MED}$ only control. Ligand is surface-coated GFP. **(b)** Drug controlled transcription via ligand mediated release of SynNotch Gal4$_{DB}$-P$_{MED}$ ICD, dNS3, and NS3 inhibitor driving UAS-H2B-mCherry expression. Ligand is surface coated c-Myc antibody. **(c)** Larger image field of view of drug-controlled ICD SynNotch cultures (see main fig. 1f for reference). Scale bar: 20 μm. Plotted values are the mean ± SD of biologically independent replicates, n=3.

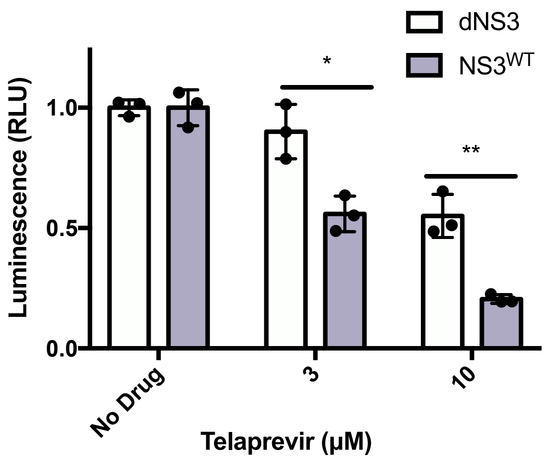


**Supplementary Figure 3 | Differential displacement profiles of dNS3 and NS3$^{WT}$ with Telaprevir.** Drug response of split dNS3-NLuc, NS3$^{WT}$-NLuc and CLuc-$P_{MED}$ with telaprevir. Dose response are normalized to their respective no drug controls. All cells were transfected 48 hours before lysing for luciferase experiments, and drug was added at the time of transfection. Plotted values are the mean ± SD of biologically independent replicates, n=3, *P<0.05, **P<0.005.

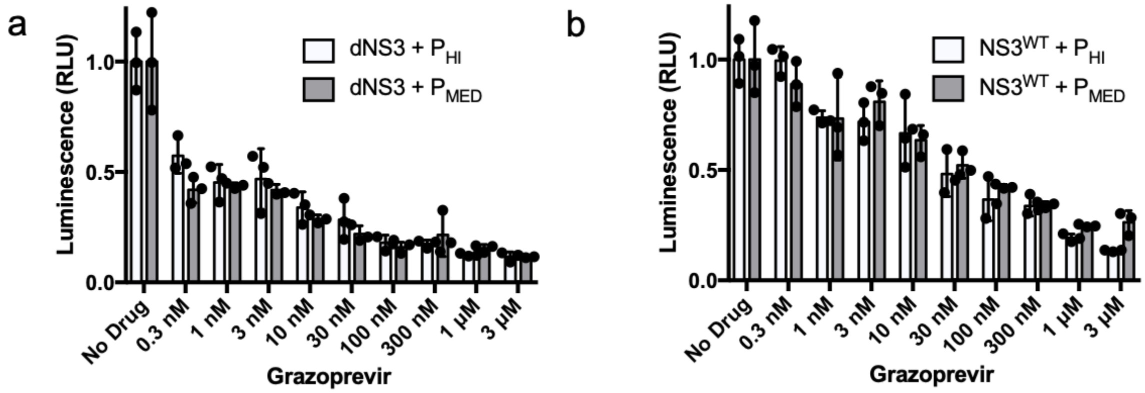


**Supplementary Figure 4 | Comparison of drug mediated displacement using $P_{MED}$ and $P_{HI}$. (a)** and **(b)** Rearranged data from main fig. 2c and 2e to compare effects of $P_{MED}$ vs. $P_{HI}$ peptides. Dose responses are normalized to their respective no drug controls. All cells were transfected 48 hours before lysing for luciferase experiments, and drug was added at the time of transfection. Plotted values are the mean ± SD of biologically independent replicates, n=3.

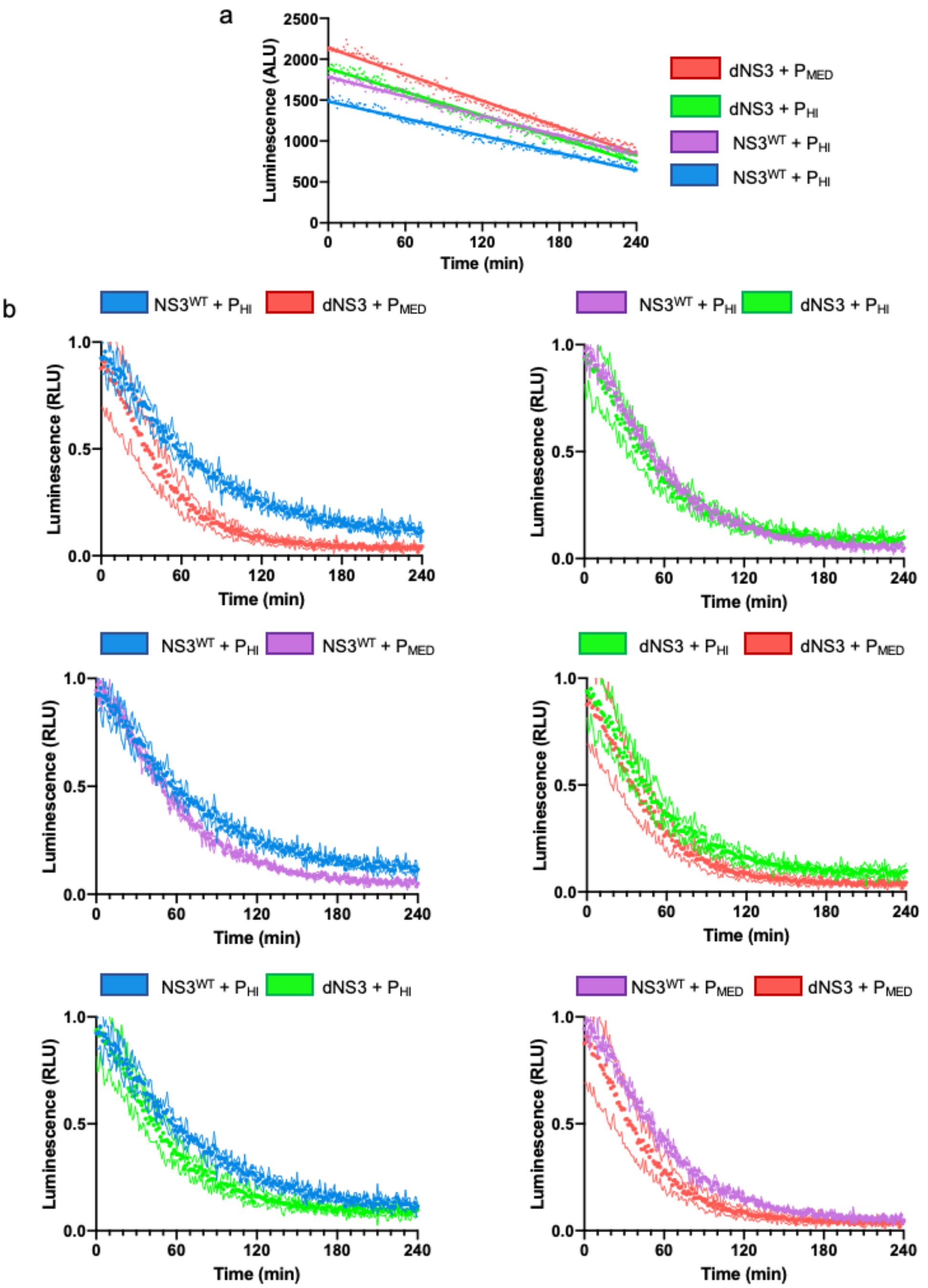


**Supplementary Figure 5 | Kinetics of displacement for various combinations of NS3$^{WT}$, dNS3, and P$_{MED}$ and P$_{HI}$ . (a)** Linear regressions of the various combinations of dNS3, NS3 and peptides to NLuc and CLuc without drug for Fig 2e. Each combination was used for normalization of each respective curve. Luminescence is not normalized here and stated in arbitrary luciferase units. **(b)** Split luciferase kinetic displacement curves rearranged from main Fig. 2e displayed as mean (dots) with SD (lines). All cells were transfected 24 hours before lysing for luciferase experiments. Experiments are biologically independent replicates, n=3. For all experiments with drug, 3 μM of grazoprevir was added at time of lysis.

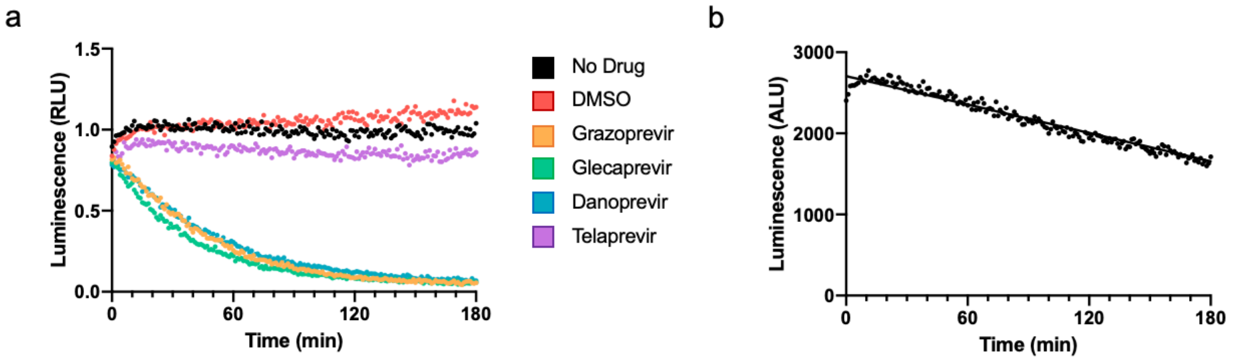


**Supplementary Figure 6 | Temporal response of dNS3-NLuc and CLuc-$P_{MED}$ to various drugs. (a)** Temporal turn-off of dNS3-NLuc and CLuc-$P_{MED}$ in response to four different antiviral NS3 inhibitors. All curves are normalized to **(b)**, a linear regression of dNS3-NLuc and CLuc-$P_{MED}$ without drug. Luminescence is not normalized here and stated in arbitrary luciferase units. All cells were transfected 24 hours before lysing for luciferase experiments. To observe temporal displacement, 3 μM of each drug was added at the time of lysis. Plotted values are the mean of biologically independent replicates, n=3.

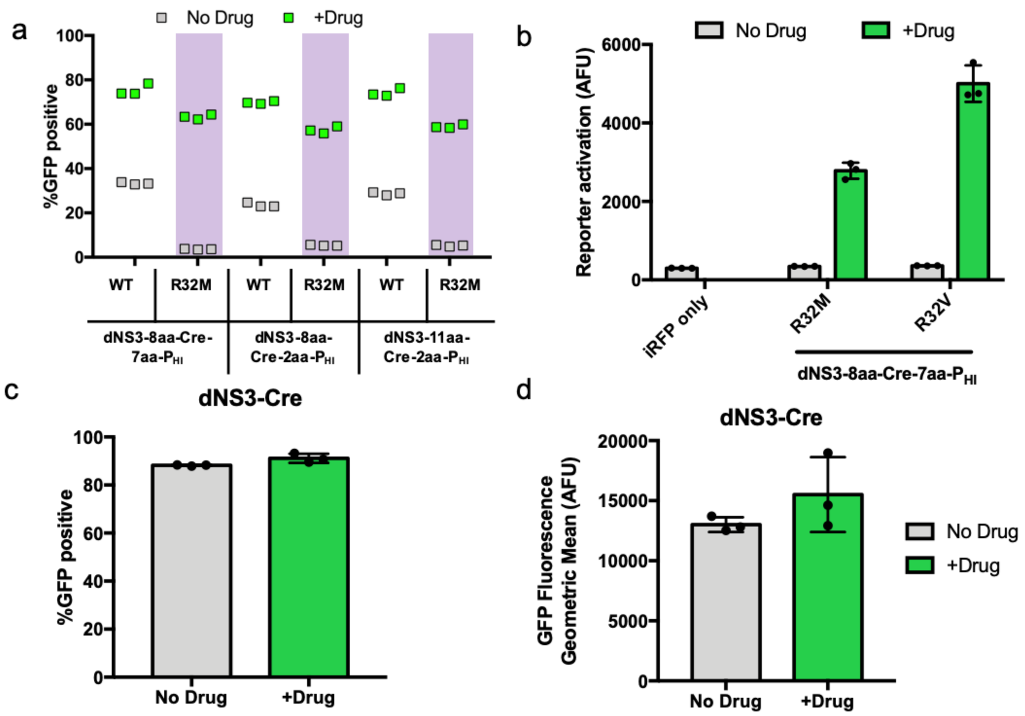


**Supplementary Figure 7| Effects of R32 mutations on various Cre constructs and geometric mean of GFP activation by Cre recombinase. (a)** Comparison of optimized linker Cre constructs with $Cre^{WT}$ and $Cre^{R32M}$ mutation. **(b)** Geometric mean fluorescent values of GFP associated with Fig. 3e. **(c)** Drug dependence of dNS3-Cre control without inhibitory peptide and **(d)** corresponding geometric mean fluorescent values of GFP, n=3 biologically independent replicates. For geometric mean fluorescence, plotted values are the geometric mean ± SD of biologically independent replicates, n=3. For all Cre constructs, drug was added 24 hours after transfection and flow cytometry data was collected 48 hours after drug addition. Drug refers 1 μM grazoprevir addition.

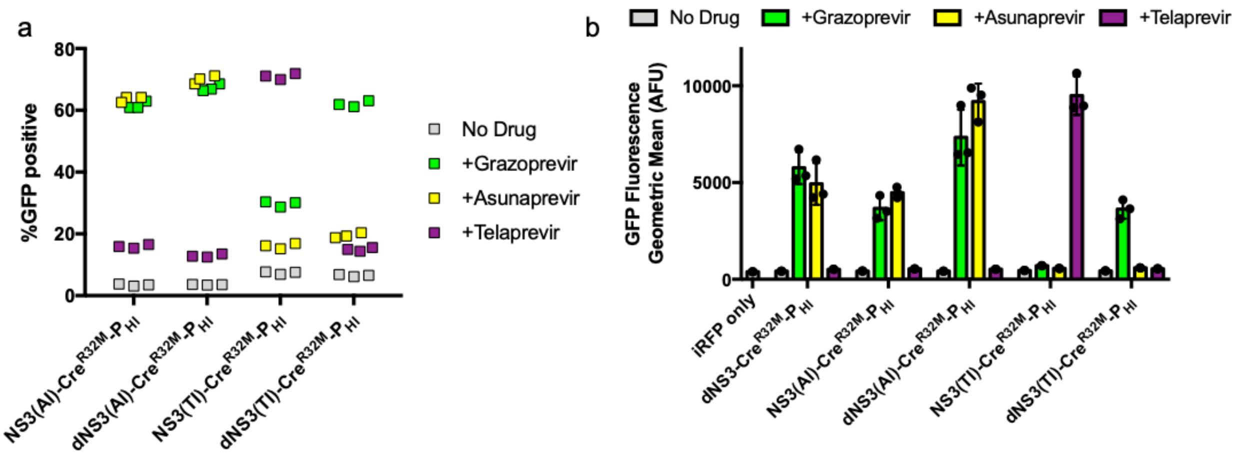


**Supplementary Figure 8| Effect of S139 mutations in drug resistant inducible Cre and geometric mean of GFP activation by drug resistant Cre recombinases. (a)** Effect of $NS3^{S139A}$ (dNS3) mutation on asunaprevir inducible (AI) and telaprevir inducible (TI) Cre recombinases. Plotted values are fraction of GFP expressing cells defined as cells with GFP signal greater than the top 1% of the non-transfected control. Basal GFP positive cells, determined by transfection marker control, were subtracted from the total, biologically independent replicates, n=3. **(b)** Associated geometric mean of fluorescence for the different drug inducible NS3 Cre recombinases with various drugs. Plotted values are the mean ± SD of biologically independent replicates, n=3. Drug was added 24 hours post-transfection, and flow cytometry data was collected 48 hours post-drug addition. Grazoprevir: 1 μM; Asunaprevir: 1 μM; Telaprevir: 10 μM.

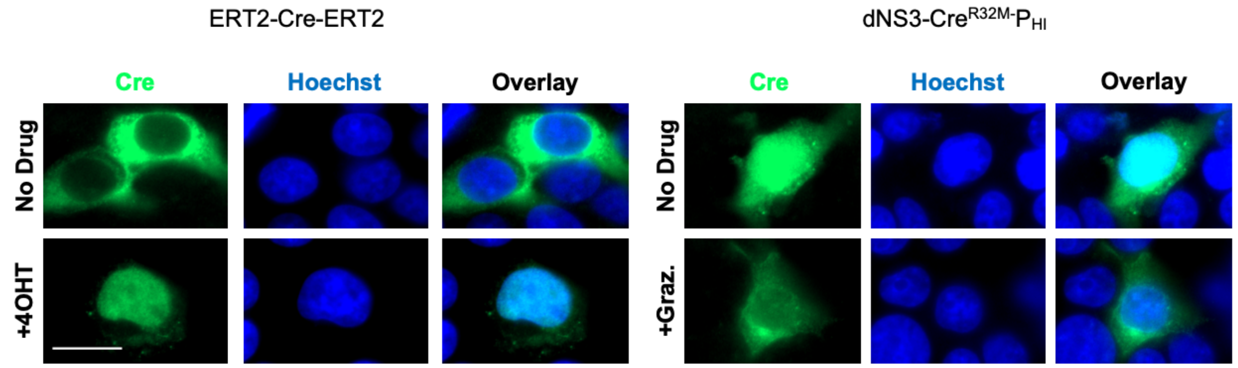


**Supplementary Figure 9| Localization of two Cre inducible systems by immunostaining.** Drug inducible Cre recombinase ERT2-Cre-ERT2 is controlled by conditional nuclear translocation upon addition of small molecule 4-hydroxytamoxifen (left). The dNS3-Cre-$P_{HI}$ constructs are present in the nucleus both with and without antiviral drug (right). Scale bar: 20 μm.

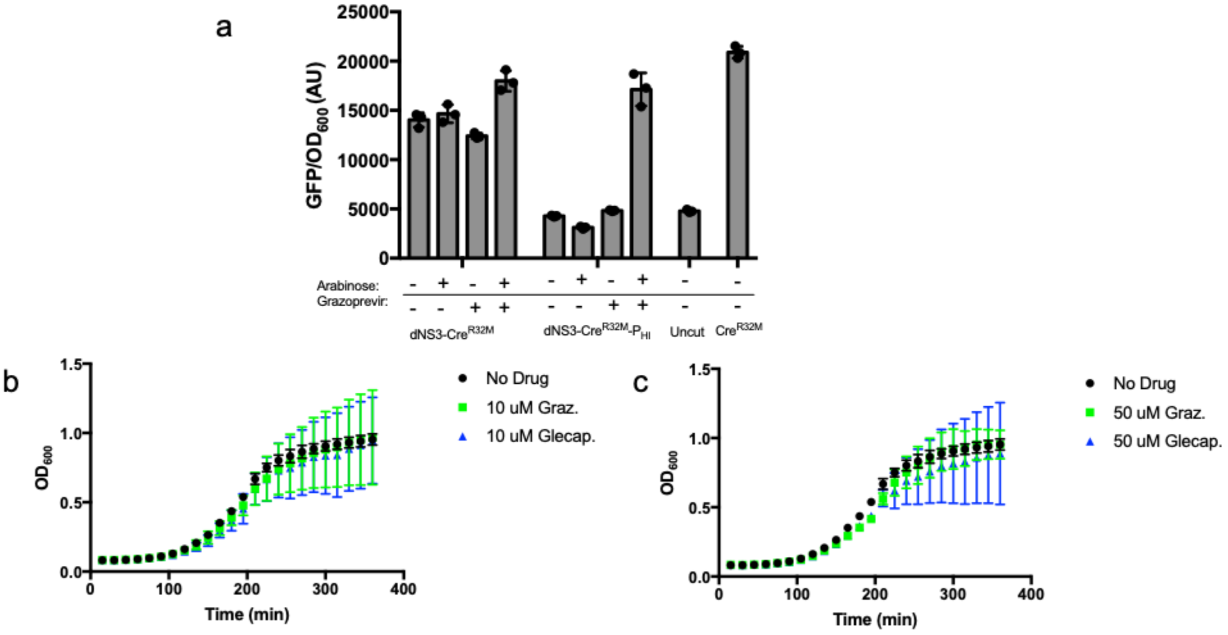


**Supplementary Figure 10| Cre reporter activation under arabinose promoters and antiviral drug effect on bacterial growth. (a)** Arabinose-inducible transcriptional control of dNS3-$Cre^{R32M}$ and dNS3-$Cre^{R32M}$ -$P_{HI}$ on low copy plasmids as compared to negative and positive control reporters, six hours post-drug addition. Arabinose: 1 mM, Grazoprevir: 50 μM **(b)** Growth curves of cells expressing dNS3-$Cre^{R32M}$ -$P_{HI}$ without drug treatment and 10 μM or **(c)** 50μM drug. Plotted values are the mean ± SD of biological replicates, n=3.

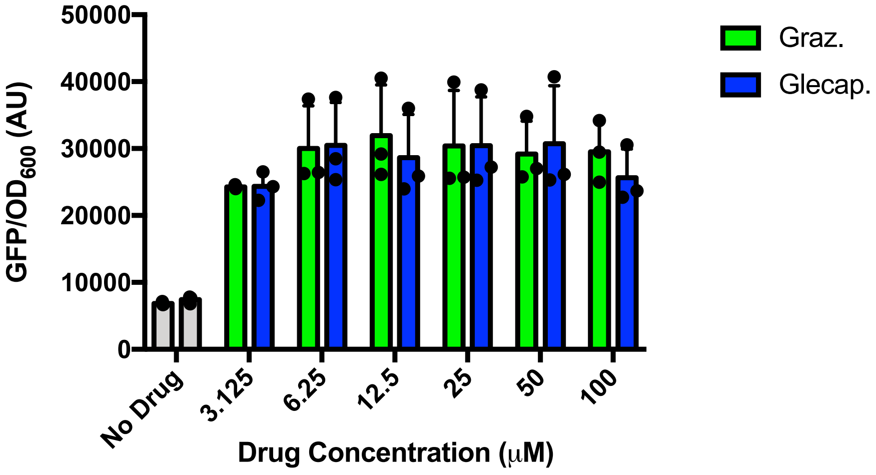


**Supplementary Figure 11 | Dose response of dNS3-Cre$^{R32M}$ -P$_{HI}$ in bacteria.** Inducible dNS3-Cre$^{R32M}$ -P$_{HI}$ was expressed under the weakest constitutive promoter in main Fig. 4b, six hours post-drug addition, fluorescence/OD quantified by plate reader. Plotted values are the mean ± SD of biological replicates, n=3.

**Supplementary Table 1 |** NS3-binding peptides used in this study.

| **Peptide** | **Pseudonym used in this study** | **Reported affinity to NS3$^{WT}$ (K$_D$)** | **Amino acid sequence** |
|---|---|---|---|
| CP5-46A-4D5E | P$_{MED}$ | 10.5 nM | GELDELVYLLDGPGYDPIHS |
| CP5-46-4D5E | P$_{HI}$ | 0.53 nM | GELDELVYLLDGPGYDPIHCDVV TRGGSHLFNF |

**Supplementary Table 2 |** Constitutive bacterial promoters used in this study in order of weakest to strongest. All sequences are derivatives of T7A1 phage promoter with mutations in the -35 and -10 regions (underlined). Transcriptional start site is in green.

| **Promoter** | **Sequence** |
|---|---|
| pW2 | TTATCAAAAAGAGTA *TTGTCT* TAAAGTCTAACCTATAG *GATTCT* TACAGCC**A**TCG AGAGGGACACGGCGAA |
| pW4 | TTATCAAAAAGAGTA *TTGCAT* TAAAGTCTAACCTATAG *GA**AT**CT* TACAGCC**A**TCG AGAGGGACACGGCGAA |
| pW6 | TTATCAAAAAGAGTA *TTGTCT* TAAAGTCTAACCTATAG GAAAAT TACAGCC**A**TCG AGAGGGACACGGCGAA |

**Supplementary Note**

Below are the relevant annotated sequences used in this work. In all sequences, NS3 and NS3 mutants contain the NS4A accessory peptide.

Gal4$_{DB}$-P$_{MED}$

ATGAAGTTGCTGAGCAGCATAGAGCAAGCATGTGATATCTGCCGGTTGAAGAAGCT
GAAGTGTAGCAAGGAGAAGCCCAAGTGCGCCAAGTGTCTCAAGAATAATTGGGAGT
GTAGGTATAGCCCCAAGACCAAGCGAAGCCCGCTTACGAGAGCACACCTTACCGAG
GTCGAGAGCCGCCTGGAAAGACTCGAACAACTTTTTCTTCTGATTTTCCCCAGGGAG
GACCTGGACATGATCCTGAAGATGGACAGCCTCCAGGACATCAAAGCCCTTCTTACC
GGGCTGTTCGTGCAGGACAACGTCAACAAGGATGCGGTGACCGACAGATTGGCGAG
CGTGGAGACGGACATGCCCTTGACCCTCAGACAACATAGGATCAGCGCGACAAGCT
CATCTGAAGAATCTAGCAATAAGGGACAGCGACAGCTGACCGTTAGTGGGGGTACC
GAATTCAGCGGGTCAGGCGGATCTGGAAGTGGTGGTGGAGAACTTGATGAATTGGT
ATACTTACTAGATGGGCCAGGTTATGACCCTATACATAGT

Gal4$_{DB}$-P$_{HI}$

ATGAAGTTGCTGAGCAGCATAGAGCAAGCATGTGATATCTGCCGGTTGAAGAAGCT
GAAGTGTAGCAAGGAGAAGCCCAAGTGCGCCAAGTGTCTCAAGAATAATTGGGAGT
GTAGGTATAGCCCCAAGACCAAGCGAAGCCCGCTTACGAGAGCACACCTTACCGAG
GTCGAGAGCCGCCTGGAAAGACTCGAACAACTTTTTCTTCTGATTTTCCCCAGGGAG
GACCTGGACATGATCCTGAAGATGGACAGCCTCCAGGACATCAAAGCCCTTCTTACC
GGGCTGTTCGTGCAGGACAACGTCAACAAGGATGCGGTGACCGACAGATTGGCGAG
CGTGGAGACGGACATGCCCTTGACCCTCAGACAACATAGGATCAGCGCGACAAGCT
CATCTGAAGAATCTAGCAATAAGGGACAGCGACAGCTGACCGTTAGTGGGGGTACC
GAATTCAGCGGGTCAGGCGGATCTGGAAGTGGTGGTGGAGAACTTGATGAATTGGT
ATACTTACTAGATGGGCCAGGTTATGACCCTATACATTGCGATGTAGTGACAAGGGG
CGGCAGCCACCTTTTCAATTTT

dNS3-VPR

S139A mutation

ATGGGAACAGGGGGTTCTGTTGTTATTGTTGGTAGAATTATTTTATCTGGTAGTGGT
AGTATCACGGCCTACTCCCAACAGACGCGGGGCCTACTTGGTTGCATCATCACTAGC
CTCACAGGCCGGGACAAGAACCAGGTCGAAGGGGAGGTTCAAGTGGTTTCTACCGC
AACACAATCTTTCCTGGCGACCTGCGTCAACGGCGTGTGCTGGACTGTCTACCATGG
CGCTGGCTCGAAGACCCTAGCCGGTCCAAAAGGTCCAATCACCCAAATGTACACCA
ATGTAGACCAGGACCTCGTCGGCTGGCAGGCGCCTCCAGGGGCGCGCTCCTTGACA
CCATGCACCTGTGGCAGCTCGGACCTTTACTTGGTCACGAGACATGCTGATGTCATT
CCGGTGCGCCGGCGAGGCGACAGCAGGGGAAGTCTACTCTCCCCCAGGCCCGTCTC
CTACCTGAAAGGCTCCGCAGGTGGTCCATTGCTTTGCCCTTCGGGGCACGCTGTGGG
CATCTTCCGGGCTGCTGTGTGCACCCGGGGGGTCGCGAAGGCGGTGGACTTCGTGCC
CGTTGAGTCTATGGAAACTACCATGCGGTCTGAGCAAAAGCTCATTTCTGAAGAGGA
CTTGTATCCCTACGATGTGCCCGATTACGCTGGCGCGTCTGCATGCCCCAAGAAGAA
GAGGAAGGTGTCGCCAGGGATCCGTCGACTTGACGCGTTGATATCAACAAGTTTGTA
CAAAAAAGCAGGCTACAAAGAGGCCAGCGGTTCCGGACGGGCTGACGCATTGGAC
GATTTTGATCTGGATATGCTGGGAAGTGACGCCCTCGATGATTTTGACCTTGACATG

CTTGGTTCGGATGCCCTTGATGACTTTGACCTCGACATGCTCGGCAGTGACGCCCTT
GATGATTTCGACCTGGACATGCTGATTAACTCTAGAAGTTCCGGATCTCCGAAAAAG
AAACGCAAAGTTGGTAGCCAGTACCTGCCCGACACCGACGACCGGCACCGGATCGA
GGAAAAGCGGAAGCGGACCTACGAGACATTCAAGAGCATCATGAAGAAGTCCCCCT
TCAGCGGCCCCACCGACCCTAGACCTCCACCTAGAAGAATCGCCGTGCCCAGCAGA
TCCAGCGCCAGCGTGCCAAAACCTGCCCCCCAGCCTTACCCCTTCACCAGCAGCCTG
AGCACCATCAACTACGACGAGTTCCCTACCATGGTGTTCCCCAGCGGCCAGATCTCT
CAGGCCTCTGCTCTGGCTCCAGCCCCTCCTCAGGTGCTGCCTCAGGCTCCTGCTCCTG
CACCAGCTCCAGCCATGGTGTCTGCACTGGCTCAGGCACCAGCACCCGTGCCTGTGC
TGGCTCCTGGACCTCCACAGGCTGTGGCTCCACCAGCCCCTAAACCTACACAGGCCG
GCGAGGGCACACTGTCTGAAGCTCTGCTGCAGCTGCAGTTCGACGACGAGGATCTG
GGAGCCCTGCTGGGAAACAGCACCGATCCTGCCGTGTTCACCGACCTGGCCAGCGT
GGACAACAGCGAGTTCCAGCAGCTGCTGAACCAGGGCATCCCTGTGGCCCCTCACA
CCACCGAGCCCATGCTGATGGAATACCCCGAGGCCATCACCCGGCTCGTGACAGGC
GCTCAGAGGCCTCCTGATCCAGCTCCTGCCCCTCTGGGAGCACCAGGCCTGCCTAAT
GGACTGCTGTCTGGCGACGAGGACTTCAGCTCTATCGCCGATATGGATTTCTCAGCC
TTGCTGGGCTCTGGCAGCGGCAGCCGGGATTCCAGGGAAGGGATGTTTTTGCCGAA
GCCTGAGGCCGGCTCCGCTATTAGTGACGTGTTTGAGGGCCGCGAGGTGTGCCAGCC
AAAACGAATCCGGCCATTTCATCCTCCAGGAAGTCCATGGGCCAACCGCCCACTCCC
CGCCAGCCTCGCACCAACACCAACCGGTCCAGTACATGAGCCAGTCGGGTCACTGA
CCCCGGCACCAGTCCCTCAGCCACTGGATCCAGCGCCCGCAGTGACTCCCGAGGCC
AGTCACCTGTTGGAGGATCCCGATGAAGAGACGAGCCAGGCTGTCAAAGCCCTTCG
GGAGATGGCCGATACTGTGATTCCCCAGAAGGAAGAGGCTGCAATCTGTGGCCAAA
TGGACCTTTCCCATCCGCCCCCAAGGGGCCATCTGGATGAGCTGACAACCACACTTG
AGTCCATGACCGAGGATCTGAACCTGGACTCACCCCTGACCCCGGAATTGAACGAG
ATTCTGGATACCTTCCTGAACGACGAGTGCCTCTTGCATGCCATGCATATCAGCACA
GGACTGTCCATCTTCGACACATCTCTGTTT

dNS3-KRAB
ATGGGAACAGGGGGTTCTGTTGTTATTGTTGGTAGAATTATTTTATCTGGTAGTGGT
AGTATCACGGCCTACTCCCAACAGACGCGGGGCCTACTTGGTTGCATCATCACTAGC
CTCACAGGCCGGGACAAGAACCAGGTCGAAGGGGAGGTTCAAGTGGTTTCTACCGC
AACACAATCTTTCCTGGCGACCTGCGTCAACGGCGTGTGCTGGACTGTCTACCATGG
CGCTGGCTCGAAGACCCTAGCCGGTCCAAAAGGTCCAATCACCCAAATGTACACCA
ATGTAGACCAGGACCTCGTCGGCTGGCAGGCGCCTCCAGGGGCGCGCTCCTTGACA
CCATGCACCTGTGGCAGCTCGGACCTTTACTTGGTCACGAGACATGCTGATGTCATT
CCGGTGCGCCGGCGAGGCGACAGCAGGGGAAGTCTACTCTCCCCCAGGCCCGTCTC
CTACCTGAAAGGCTCCGCAGGTGGTCCATTGCTTTGCCCTTCGGGGCACGCTGTGGG
CATCTTCCGGGCTGCTGTGTGCACCCGGGGGGTCGCGAAGGCGGTGGACTTCGTGCC
CGTTGAGTCTATGGAAACTACCATGCGGTCTGAGCAAAAGCTCATTTCTGAAGAGGA
CTTGTATCCCTACGATGTGCCCGATTACGCTGGCGCGTCTGCATGCCCCAAGAAGAA
GAGGAAGGTGTCGCCAGGGATCCGTCGACTTGACGCGTTGATATCAACAAGTTTGTA
CAAAAAAGCAGGCTACAAAGAGGCCAGCGGTTCCACCGGTATGGATGCTAAGTCAC
TAACTGCCTGGTCCCGGACACTGGTGACCTTCAAGGATGTATTTGTGGACTTCACCA
GGGAGGAGTGGAAGCTGCTGGACACTGCTCAGCAGATCGTGTACAGAAATGTGATG

CTGGAGAACTATAAGAACCTGGTTTCCTTGGGTTATCAGCTTACTAAGCCAGATGTG
ATCCTCCGGTTGGAGAAGGGAGAAGAGCCC

SS-Anti-GFP-SynNotch-Gal4$_{DB}$-P$_{MED}$
ATGGCATTGCCCGTGACCGCCCTGCTGCTGCCACTGGCCTTGTTGCTCCACGCCGCG
CGGCCAGAACAGAAGCTGATCAGCGAGGAGGATCTGATGGCTGATGTGCAACTCGT
AGAGTCCGGTGGCGGACTCGTCCAGGCGGGAGGCTCCCTGCGCCTGAGTTGCGCTG
CGTCCGGAAGAACTATTAGCATGGCCGCCATGAGTTGGTTCAGACAGGCGCCTGGA
AAGGAGAGGGAGTTCGTGGCAGGCATAAGTCGCTCCGCAGGGTCAGCCGTCCATGC
CGACTCCGTAAAGGGACGCTTTACCATCTCCAGAGATAACACAAAAAACACGCTCT
ATCTCCAAATGAATTCTTTGAAGGCCGAGGACACAGCCGTGTACTACTGCGCCGTTA
GGACCAGCGGATTTTTTGGAAGCATTCCTAGGACAGGGACAGCCTTTGATTACTGGG
GCCAGGGAACGCAGGTTACCGTTTCCATCCTGGACTACAGCTTCACAGGTGGCGCTG
GGCGCGACATTCCCCCACCGCAGATTGAGGAGGCCTGTGAGCTGCCTGAGTGCCAG
GTGGATGCAGGCAATAAGGTCTGCAACCTGCAGTGTAATAATCACGCATGTGGCTG
GGATGGTGGCGACTGCTCCCTCAACTTCAATGACCCCTGGAAGAACTGCACGCAGTC
TCTACAGTGCTGGAAGTATTTTAGCGACGGCCACTGTGACAGCCAGTGCAACTCGGC
CGGCTGCCTCTTTGATGGCTTCGACTGCCAGCTCACCGAGGGACAGTGCAACCCCCT
GTATGACCAGTACTGCAAGGACCACTTCAGTGATGGCCACTGCGACCAGGGCTGTA
ACAGTGCCGAATGTGAGTGGGATGGCCTAGACTGTGCTGAGCATGTACCCGAGCGG
CTGGCAGCCGGCACCCTGGTGCTGGTGGTGCTGCTTCCACCCGACCAGCTACGGAAC
AACTCCTTCCACTTTCTGCGGGAGCTCAGCCACGTGCTGCACACCAACGTGGTCTTC
AAGCGTGATGCGCAAGGCCAGCAGATGATCTTCCCGTACTATGGCCACGAGGAAGA
GCTGCGCAAGCACCCAATCAAGCGCTCTACAGTGGGTTGGGCCACCTCTTCACTGCT
TCCTGGTACCAGTGGTGGGCGCCAGCGCAGGGAGCTGGACCCCATGGACATCCGTG
GCTCCATTGTCTACCTGGAGATCGACAACCGGCAATGTGTGCAGTCATCCTCGCAGT
GCTTCCAGAGTGCCACCGATGTGGCTGCCTTCCTAGGTGCTCTTGCGTCACTTGGCA
GCCTCAATATTCCTTACAAGATTGAGGCCGTGAAGAGTGAGCCGGTGGAGCCTCCGC
TGCCCTCGCAGCTGCACCTCATGTACGTGGCAGCGGCCGCCTTCGTGCTCCTGTTCTT
TGTGGGCTGTGGGGTGCTGCTGTCCCGCAAGCGCCGGCGGATGAAGCTGCTGAGCA
GCATAGAGCAAGCATGTGATATCTGCCGGTTGAAGAAGCTGAAGTGTAGCAAGGAG
AAGCCCAAGTGCGCCAAGTGTCTCAAGAATAATTGGGAGTGTAGGTATAGCCCCAA
GACCAAGCGAAGCCCGCTTACGAGAGCACACCTTACCGAGGTCGAGAGCCGCCTGG
AAAGACTCGAACAACTTTTTCTTCTGATTTTCCCCAGGGAGGACCTGGACATGATCC
TGAAGATGGACAGCCTCCAGGACATCAAAGCCCTTCTTACCGGGCTGTTCGTGCAGG
ACAACGTCAACAAGGATGCGGTGACCGACAGATTGGCGAGCGTGGAGACGGACATG
CCCTTGACCCTCAGACAACATAGGATCAGCGCGACAAGCTCATCTGAAGAATCTAG
CAATAAGGGACAGCGACAGCTGACCGTTAGTGGGGGTACCGAATTCAGCGGGTCAG
GCGGATCTGGAAGTGGTGGTGGAGAACTTGATGAATTGGTATACTTACTAGATGGG
CCAGGTTATGACCCTATACATAGT

dNS3-NLuc

ATGGGAACAGGGGGTTCTGTTGTTATTGTTGGTAGAATTATTTTATCTGGTAGTGGT
AGTATCACGGCCTACTCCCAACAGACGCGGGGCCTACTTGGTTGCATCATCACTAGC
CTCACAGGCCGGGACAAGAACCAGGTCGAAGGGGAGGTTCAAGTGGTTTCTACCGC
AACACAATCTTTCCTGGCGACCTGCGTCAACGGCGTGTGCTGGACTGTCTACCATGG
CGCTGGCTCGAAGACCCTAGCCGGTCCAAAAGGTCCAATCACCCAAATGTACACCA
ATGTAGACCAGGACCTCGTCGGCTGGCAGGCGCCTCCAGGGGCGCGCTCCTTGACA
CCATGCACCTGTGGCAGCTCGGACCTTTACTTGGTCACGAGACATGCTGATGTCATT
CCGGTGCGCCGGCGAGGCGACAGCAGGGGAAGTCTACTCTCCCCCAGGCCCGTCTC
CTACCTGAAAGGCTCCGCAGGTGGTCCATTGCTTTGCCCTTCGGGGCACGCTGTGGG
CATCTTCCGGGCTGCTGTGTGCACCCGGGGGGTCGCGAAGGCGGTGGACTTCGTGCC
CGTTGAGTCTATGGAAACTACCATGCGGTCTTCGTACGCGTCCCGGGGCGGTGGCTC
ATCTGGCGGAGGTGAAGACGCCAAAAACATAAAGAAAGGCCCGGCGCCATTCTATC
CGCTGGAAGATGGAACCGCTGGAGAGCAACTGCATAAGGCTATGAAGAGATACGCC
CTGGTTCCTGGAACAATTGCTTTTACAGATGCACATATCGAGGTGGACATCACTTAC
GCTGAGTACTTCGAAATGTCCGTTCGGTTGGCAGAAGCTATGAAACGATATGGGCTG
AATACAAATCACAGAATCGTCGTATGCAGTGAAAACTCTCTTCAATTCTTTATGCCG
GTGTTGGGCGCGTTATTTATCGGAGTTGCAGTTGCGCCCGCGAACGACATTTATAAT
GAACGTGAATTGCTCAACAGTATGGGCATTTCGCAGCCTACCGTGGTGTTCGTTTCC
AAAAAGGGGTTGCAAAAAATTTTGAACGTGCAAAAAAAGCTCCCAATCATCCAAAA
AATTATTATCATGGATTCTAAAACGGATTACCAGGGATTTCAGTCGATGTACACGTT
CGTCACATCTCATCTACCTCCCGGTTTTAATGAATACGATTTTGTGCCAGAGTCCTTC
GATAGGGACAAGACAATTGCACTGATCATGAACTCCTCTGGATCTACTGGTCTGCCT
AAAGGTGTCGCTCTGCCTCATAGAACTGCCTGCGTGAGATTCTCGCATGCCAGAGAT
CCTATTTTTGGCAATCAAATCATTCCGGATACTGCGATTTTAAGTGTTGTTCCATTCC
ATCACGGTTTTGGAATGTTTACTACACTCGGATATTTGATATGTGGATTTCGAGTCGT
CTTAATGTATAGATTTGAAGAAGAGCTGTTTCTGAGGAGCCTTCAGGATTACAAGAT
TCAAAGTGCGCTGCTGGTGCCAACCCTATTCTCCTTCTTCGCCAAAAGCACTCTGATT
GACAAATACGATTTATCTAATTTACACGAAATTGCTTCTGGTGGCGCTCCCCTCTCTA
AGGAAGTCGGGGAAGCGGTTGCCAAGAGGTTCCATCTGCCAGGTATCAGGCAAGGA
TATGGGCTCACTGAGACTACATCAGCTATTCTGATTACACCCGAGGGGGATGATAAA
CCGGGCGCGGTCGGTAAAGTTGTTCCATTTTTTGAAGCGAAGGTTGTGGATCTGGAT
ACCGGGAAAACGCTGGGCGTTAATCAAAGAGGCGAACTGTGTGTGAGAGGTCCTAT
GATTATGTCCGGTTATGTAAACAATCCGGAAGCGACCAACGCCTTGATTGACAAGG
ATGGA

NS3$^{WT}$-NLuc

ATGGGAACAGGGGGTTCTGTTGTTATTGTTGGTAGAATTATTTTATCTGGTAGTGGT
AGTATCACGGCCTACTCCCAACAGACGCGGGGCCTACTTGGTTGCATCATCACTAGC
CTCACAGGCCGGGACAAGAACCAGGTCGAAGGGGAGGTTCAAGTGGTTTCTACCGC
AACACAATCTTTCCTGGCGACCTGCGTCAACGGCGTGTGCTGGACTGTCTACCATGG
CGCTGGCTCGAAGACCCTAGCCGGTCCAAAAGGTCCAATCACCCAAATGTACACCA
ATGTAGACCAGGACCTCGTCGGCTGGCAGGCGCCTCCAGGGGCGCGCTCCTTGACA
CCATGCACCTGTGGCAGCTCGGACCTTTACTTGGTCACGAGACATGCTGATGTCATT
CCGGTGCGCCGGCGAGGCGACAGCAGGGGAAGTCTACTCTCCCCCAGGCCCGTCTC
CTACCTGAAAGGCTCCTCAGGTGGTCCATTGCTTTGCCCTTCGGGGCACGCTGTGGG

CATCTTCCGGGCTGCTGTGTGCACCCGGGGGGTCGCGAAGGCGGTGGACTTCGTGCC
CGTTGAGTCTATGGAAACTACCATGCGGTCTTCGTACGCGTCCCGGGGCGGTGGCTC
ATCTGGCGGAGGTGAAGACGCCAAAAACATAAAGAAAGGCCCGGCGCCATTCTATC
CGCTGGAAGATGGAACCGCTGGAGAGCAACTGCATAAGGCTATGAAGAGATACGCC
CTGGTTCCTGGAACAATTGCTTTTACAGATGCACATATCGAGGTGGACATCACTTAC
GCTGAGTACTTCGAAATGTCCGTTCGGTTGGCAGAAGCTATGAAACGATATGGGCTG
AATACAAATCACAGAATCGTCGTATGCAGTGAAAACTCTCTTCAATTCTTTATGCCG
GTGTTGGGCGCGTTATTTATCGGAGTTGCAGTTGCGCCCGCGAACGACATTTATAAT
GAACGTGAATTGCTCAACAGTATGGGCATTTCGCAGCCTACCGTGGTGTTCGTTTCC
AAAAAGGGGTTGCAAAAAATTTTGAACGTGCAAAAAAAGCTCCCAATCATCCAAAA
AATTATTATCATGGATTCTAAAACGGATTACCAGGGATTTCAGTCGATGTACACGTT
CGTCACATCTCATCTACCTCCCGGTTTTAATGAATACGATTTTGTGCCAGAGTCCTTC
GATAGGGACAAGACAATTGCACTGATCATGAACTCCTCTGGATCTACTGGTCTGCCT
AAAGGTGTCGCTCTGCCTCATAGAACTGCCTGCGTGAGATTCTCGCATGCCAGAGAT
CCTATTTTTGGCAATCAAATCATTCCGGATACTGCGATTTTAAGTGTTGTTCCATTCC
ATCACGGTTTTGGAATGTTTACTACACTCGGATATTTGATATGTGGATTTCGAGTCGT
CTTAATGTATAGATTTGAAGAAGAGCTGTTTCTGAGGAGCCTTCAGGATTACAAGAT
TCAAAGTGCGCTGCTGGTGCCAACCCTATTCTCCTTCTTCGCCAAAAGCACTCTGATT
GACAAATACGATTTATCTAATTTACACGAAATTGCTTCTGGTGGCGCTCCCCTCTCTA
AGGAAGTCGGGGAAGCGGTTGCCAAGAGGTTCCATCTGCCAGGTATCAGGCAAGGA
TATGGGCTCACTGAGACTACATCAGCTATTCTGATTACACCCGAGGGGGATGATAAA
CCGGGCGCGGTCGGTAAAGTTGTTCCATTTTTTGAAGCGAAGGTTGTGGATCTGGAT
ACCGGGAAAACGCTGGGCGTTAATCAAAGAGGCGAACTGTGTGTGAGAGGTCCTAT
GATTATGTCCGGTTATGTAAACAATCCGGAAGCGACCAACGCCTTGATTGACAAGG
ATGGA

CLuc-$P_{MED}$

ATGTCCGGTTATGTAAACAATCCGGAAGCGACCAACGCCTTGATTGACAAGGATGG
ATGGCTACATTCTGGAGACATAGCTTACTGGGACGAAGACGAACACTTCTTCATCGT
TGACCGCCTGAAGTCTCTGATTAAGTACAAAGGCTATCAGGTGGCTCCCGCTGAATT
GGAATCCATCTTGCTCCAACACCCCAACATCTTCGACGCAGGTGTCGCAGGTCTTCC
CGACGATGACGCCGGTGAACTTCCCGCCGCCGTTGTTGTTTTGGAGCACGGAAAGAC
GATGACGGAAAAAGAGATCGTGGATTACGTCGCCAGTCAAGTAACAACCGCGAAAA
AGTTGCGCGGAGGAGTTGTGTTTGTGGACGAAGTACCGAAAGGTCTTACCGGAAAA
CTCGACGCAAGAAAAATCAGAGAGATCCTCATAAAGGCCAAGAAGGGCGGAAAGA
TCGCCGTGGGAGGTGGCTCATCTGGCGGAGGTCAGATCTCGTACGGTGGAGAACTT
GATGAATTGGTATACTTACTAGATGGGCCAGGTTATGACCCTATACATAGT

CLuc-$P_{HI}$

ATGTCCGGTTATGTAAACAATCCGGAAGCGACCAACGCCTTGATTGACAAGGATGG
ATGGCTACATTCTGGAGACATAGCTTACTGGGACGAAGACGAACACTTCTTCATCGT
TGACCGCCTGAAGTCTCTGATTAAGTACAAAGGCTATCAGGTGGCTCCCGCTGAATT
GGAATCCATCTTGCTCCAACACCCCAACATCTTCGACGCAGGTGTCGCAGGTCTTCC
CGACGATGACGCCGGTGAACTTCCCGCCGCCGTTGTTGTTTTGGAGCACGGAAAGAC

GATGACGGAAAAAGAGATCGTGGATTACGTCGCCAGTCAAGTAACAACCGCGAAAA
AGTTGCGCGGAGGAGTTGTGTTTGTGGACGAAGTACCGAAAGGTCTTACCGGAAAA
CTCGACGCAAGAAAAATCAGAGAGATCCTCATAAAGGCCAAGAAGGGCGGAAAGA
TCGCCGTGGGAGGTGGCTCATCTGGCGGAGGTCAGATCTCGTACGGTGGAGAACTT
GATGAATTGGTATACTTACTAGATGGGCCAGGTTATGACCCTATACATTGCGATGTA
GTGACAAGGGGCGGCAGCCACCTTTTCAATTTT

dNS3-8aa-Cre$^{R32M}$-7aa-P$_{HI}$

ATGGGAACAGGGGGTTCTGTTGTTATTGTTGGTAGAATTATTTATCTGGTAGTGGT
AGTATCACGGCCTACTCCCAACAGACGCGGGGCCTACTTGGTTGCATCATCACTAGC
CTCACAGGCCGGGACAAGAACCAGGTCGAAGGGGAGGTTCAAGTGGTTTCTACCGC
AACACAATCTTTCCTGGCGACCTGCGTCAACGGCGTGTGCTGGACTGTCTACCATGG
CGCTGGCTCGAAGACCCTAGCCGGTCCAAAAGGTCCAATCACCCAAATGTACACCA
ATGTAGACCAGGACCTCGTCGGCTGGCAGGCGCCTCCAGGGGCGCGCTCCTTGACA
CCATGCACCTGTGGCAGCTCGGACCTTTACTTGGTCACGAGACATGCTGATGTCATT
CCGGTGCGCCGGCGAGGCGACAGCAGGGGAAGTCTACTCTCCCCCAGGCCCGTCTC
CTACCTGAAAGGCTCCGCAGGTGGTCCATTGCTTTGCCCTTCGGGGCACGCTGTGGG
CATCTTCCGGGCTGCTGTGTGCACCCGGGGGGTCGCGAAGGCGGTGGACTTCGTGCC
CGTTGAGTCTATGGAAACTACCATGCGGTCTGGTACCAGTGGCGGTAGTCAATTGAA
TTTACTGACCGTACACCAAAATTTGCCTGCATTACCGGTCGATGCAACGAGTGATGA
GGTTCGCAAGAACCTGATGGACATGTTCATGGATCGCCAGGCGTTTTCTGAGCATAC
CTGGAAAATGCTTCTGTCCGTTTGCCGGTCGTGGGCGGCATGGTGCAAGTTGAATAA
CCGGAAATGGTTTCCCGCAGAACCTGAAGATGTTCGCGATTATCTTCTATATCTTCA
GGCGCGCGGTCTGGCAGTAAAAACTATCCAGCAACATTTGGGCCAGCTAAACATGC
TTCATCGTCGGTCCGGGCTGCCACGACCAAGTGACAGCAATGCTGTTTCACTGGTTA
TGCGGCGGATCCGAAAAGAAAACGTTGATGCCGGTGAACGTGCAAAACAGGCTCTA
GCGTTCGAACGCACTGATTTCGACCAGGTTCGTTCACTCATGGAAAATAGCGATCGC
TGCCAGGATATACGTAATCTGGCATTTCTGGGGATTGCTTATAACACCCTGTTACGT
ATAGCCGAAATTGCCAGGATCAGGGTTAAAGATATCTCACGTACTGACGGTGGGAG
AATGTTAATCCATATTGGCAGAACGAAAACGCTGGTTAGCACCGCAGGTGTAGAGA
AGGCACTTAGCCTGGGGGTAACTAAACTGGTCGAGCGATGGATTTCCGTCTCTGGTG
TAGCTGATGATCCGAATAACTACCTGTTTTGCCGGGTCAGAAAAAATGGTGTTGCCG
CGCCATCTGCCACCAGCCAGCTATCAACTCGCGCCCTGGAAGGGATTTTTGAAGCAA
CTCATCGATTGATTTACGGCGCTAAGGATGACTCTGGTCAGAGATACCTGGCCTGGT
CTGGACACAGTGCCCGTGTCGGAGCCGCGCGAGATATGGCCCGCGCTGGAGTTTCA
ATACCGGAGATCATGCAAGCTGGTGGCTGGACCAATGTAAATATTGTCATGAACTAT
ATCCGTAACCTGGATAGTGAAACAGGGGCAATGGTGCGCCTGCTGGAAGATGGCGA
TCTCGAGTCTGGAAGTGGTGGTGGAGAACTTGATGAATTGGTATACTTACTAGATGG
GCCAGGTTATGACCCTATACATTGCGATGTAGTGACAAGGGGCGGCAGCCACCTTTT
CAATTTT

dNS3-8aa-Cre$^{R32V}$-7aa-P$_{HI}$

ATGGGAACAGGGGGTTCTGTTGTTATTGTTGGTAGAATTATTTATCTGGTAGTGGT
AGTATCACGGCCTACTCCCAACAGACGCGGGGCCTACTTGGTTGCATCATCACTAGC
CTCACAGGCCGGGACAAGAACCAGGTCGAAGGGGAGGTTCAAGTGGTTTCTACCGC
AACACAATCTTTCCTGGCGACCTGCGTCAACGGCGTGTGCTGGACTGTCTACCATGG

CGCTGGCTCGAAGACCCTAGCCGGTCCAAAAGGTCCAATCACCCAAATGTACACCA
ATGTAGACCAGGACCTCGTCGGCTGGCAGGCGCCTCCAGGGGCGCGCTCCTTGACA
CCATGCACCTGTGGCAGCTCGGACCTTTACTTGGTCACGAGACATGCTGATGTCATT
CCGGTGCGCCGGCGAGGCGACAGCAGGGGAAGTCTACTCTCCCCCAGGCCCGTCTC
CTACCTGAAAGGCTCCGCAGGTGGTCCATTGCTTTGCCCTTCGGGGCACGCTGTGGG
CATCTTCCGGGCTGCTGTGTGCACCCGGGGGGTCGCGAAGGCGGTGGACTTCGTGCC
CGTTGAGTCTATGGAAACTACCATGCGGTCTGGTACCAGTGGCGGTAGTCAATTGAA
TTTACTGACCGTACACCAAAATTTGCCTGCATTACCGGTCGATGCAACGAGTGATGA
GGTTCGCAAGAACCTGATGGACATGTTCGTGGATCGCCAGGCGTTTTCTGAGCATAC
CTGGAAAATGCTTCTGTCCGTTTGCCGGTCGTGGGCGGCATGGTGCAAGTTGAATAA
CCGGAAATGGTTTCCCGCAGAACCTGAAGATGTTCGCGATTATCTTCTATATCTTCA
GGCGCGCGGTCTGGCAGTAAAAACTATCCAGCAACATTTGGGCCAGCTAAACATGC
TTCATCGTCGGTCCGGGCTGCCACGACCAAGTGACAGCAATGCTGTTTCACTGGTTA
TGCGGCGGATCCGAAAAGAAAACGTTGATGCCGGTGAACGTGCAAAACAGGCTCTA
GCGTTCGAACGCACTGATTTCGACCAGGTTCGTTCACTCATGGAAAATAGCGATCGC
TGCCAGGATATACGTAATCTGGCATTTCTGGGGATTGCTTATAACACCCTGTTACGT
ATAGCCGAAATTGCCAGGATCAGGGTTAAAGATATCTCACGTACTGACGGTGGGAG
AATGTTAATCCATATTGGCAGAACGAAAACGCTGGTTAGCACCGCAGGTGTAGAGA
AGGCACTTAGCCTGGGGGTAACTAAACTGGTCGAGCGATGGATTTCCGTCTCTGGTG
TAGCTGATGATCCGAATAACTACCTGTTTTGCCGGGTCAGAAAAAATGGTGTTGCCG
CGCCATCTGCCACCAGCCAGCTATCAACTCGCGCCCTGGAAGGGATTTTTGAAGCAA
CTCATCGATTGATTTACGGCGCTAAGGATGACTCTGGTCAGAGATACCTGGCCTGGT
CTGGACACAGTGCCCGTGTCGGAGCCGCGCGAGATATGGCCCGCGCTGGAGTTTCA
ATACCGGAGATCATGCAAGCTGGTGGCTGGACCAATGTAAATATTGTCATGAACTAT
ATCCGTAACCTGGATAGTGAAACAGGGGCAATGGTGCGCCTGCTGGAAGATGGCGA
TCTCGAGTCTGGAAGTGGTGGTGGAGAACTTGATGAATTGGTATACTTACTAGATGG
GCCAGGTTATGACCCTATACATTGCGATGTAGTGACAAGGGGCGGCAGCCACCTTTT
CAATTTT

dNS3(AI)-8aa-Cre$^{R32M}$-7aa-P$_{HI}$

ATGGGAACAGGGGGTTCTGTTGTTATTGTTGGTAGAATTATTTATCTGGTAGTGGT
AGTATCACGGCCTACTCCCAACAGACGCGGGGCCTACTTGGTTGCATCATCACTAGC
CTCACAGGCCGGGACAAGAACCAGGTCGAAGGGGAGGTTCAAGTGATGTCTACCGC
AACACAATCTTTCCTGGCGACCTGCGTCAACGGCGTGTGCTGGGCGGTCTACCATGG
CGCTGGCTCGAAGACCCTAGCCGGTCCAAAAGGTCCAATCACCCAAATGTACACCA
ATGTAGACCAGGACCTCGTCGGCTGGCAGGCGCCTCCAGGGGCGCGCTCCTTGACA
CCATGCACCTGTGGCAGCTCGGACCTTTACTTGGTCACGAGACATGCTGATGTCATT
CCGGTGCGCCGGCGAGGCGACGGGAGGGGAAGTCTACTCTCCCCCAGGCCCGTCTC
CTACCTGAAAGGCTCCGCAGGTGGTCCATTGCTTTGCCCTTCGGGGCACGCTGTGGG
CATCTTCCGGGCTGCTGTGTGCACCCGGGGGGTCGCGAAGGCGGTGGACTTCGTGCC
CGTTGAGTCTATGGAAACTACCATGCGGTCTGGTACCAGTGGCGGTAGTCAATTGAA
TTTACTGACCGTACACCAAAATTTGCCTGCATTACCGGTCGATGCAACGAGTGATGA
GGTTCGCAAGAACCTGATGGACATGTTCATGGATCGCCAGGCGTTTTCTGAGCATAC
CTGGAAAATGCTTCTGTCCGTTTGCCGGTCGTGGGCGGCATGGTGCAAGTTGAATAA
CCGGAAATGGTTTCCCGCAGAACCTGAAGATGTTCGCGATTATCTTCTATATCTTCA

GGCGCGCGGTCTGGCAGTAAAAACTATCCAGCAACATTTGGGCCAGCTAAACATGC
TTCATCGTCGGTCCGGGCTGCCACGACCAAGTGACAGCAATGCTGTTTCACTGGTTA
TGCGGCGGATCCGAAAAGAAAACGTTGATGCCGGTGAACGTGCAAAACAGGCTCTA
GCGTTCGAACGCACTGATTTCGACCAGGTTCGTTCACTCATGGAAAATAGCGATCGC
TGCCAGGATATACGTAATCTGGCATTTCTGGGGATTGCTTATAACACCCTGTTACGT
ATAGCCGAAATTGCCAGGATCAGGGTTAAAGATATCTCACGTACTGACGGTGGGAG
AATGTTAATCCATATTGGCAGAACGAAAACGCTGGTTAGCACCGCAGGTGTAGAGA
AGGCACTTAGCCTGGGGGTAACTAAACTGGTCGAGCGATGGATTTCCGTCTCTGGTG
TAGCTGATGATCCGAATAACTACCTGTTTTGCCGGGTCAGAAAAAATGGTGTTGCCG
CGCCATCTGCCACCAGCCAGCTATCAACTCGCGCCCTGGAAGGGATTTTTGAAGCAA
CTCATCGATTGATTTACGGCGCTAAGGATGACTCTGGTCAGAGATACCTGGCCTGGT
CTGGACACAGTGCCCGTGTCGGAGCCGCGCGAGATATGGCCCGCGCTGGAGTTTCA
ATACCGGAGATCATGCAAGCTGGTGGCTGGACCAATGTAAATATTGTCATGAACTAT
ATCCGTAACCTGGATAGTGAAACAGGGGCAATGGTGCGCCTGCTGGAAGATGGCGA
TCTCGAGTCTGGAAGTGGTGGTGGAGAACTTGATGAATTGGTATACTTACTAGATGG
GCCAGGTTATGACCCTATACATTGCGATGTAGTGACAAGGGGCGGCAGCCACCTTTT
CAATTTT

NS3(TI)-8aa-Cre$^{R32M}$-7aa-P$_{HI}$
Note: No S139A mutation
ATGGGAACAGGGGGTTCTGTTGTTATTGTTGGTAGAATTATTTTATCTGGTAGTGGT
AGTATCACGGCCTACTCCCAACAGACGCGGGGCCTACTTGGTTGCATCATCACTAGC
CTCACAGGCCGGGACAAGAACCAGGTCGAAGGGGAGGTTCAAGTGGTTTCTACCGC
AACACAATCTCTTCTGGCGACCTGCGTCAACGGCGTGTGCTGGACTGTCTACCATGG
CGCTGGCTCGAAGACCCTAGCCGGTCCAAAAGGTCCAATCACCCAAATGTACACCA
ATGTAGACAAAGACCTCGTCGGCTGGCAGGCGCCTCCAGGGGCGCGCTCCTTGACA
CCATGCACCTGTGGCAGCTCGGACCTTTACTTGGTCACGAGACATGCTGATGTCATT
CCGGTGCGCCGGCGAGGCGACAGGAGGGGAAGTCTACTCTCCCCCAGGCCCGTCTC
CTACCTGAAAGGCTCCTCAGGTGGTCCATTGCTTTGCCCTTCGGGGCACGCTGTGGG
CATCTTCCGGGCTGCTGTGTGCACCCGGGGGGTCGCGAAGGCGGTGTACTTCGTGCC
CGTTGAGTCTATGGAAACTACCATGCGGTCTGGTACCAGTGGCGGTAGTCAATTGAA
TTTACTGACCGTACACCAAAATTTGCCTGCATTACCGGTCGATGCAACGAGTGATGA
GGTTCGCAAGAACCTGATGGACATGTTCATGGATCGCCAGGCGTTTTCTGAGCATAC
CTGGAAAATGCTTCTGTCCGTTTGCCGGTCGTGGGCGGCATGGTGCAAGTTGAATAA
CCGGAAATGGTTTCCCGCAGAACCTGAAGATGTTCGCGATTATCTTCTATATCTTCA
GGCGCGCGGTCTGGCAGTAAAAACTATCCAGCAACATTTGGGCCAGCTAAACATGC
TTCATCGTCGGTCCGGGCTGCCACGACCAAGTGACAGCAATGCTGTTTCACTGGTTA
TGCGGCGGATCCGAAAAGAAAACGTTGATGCCGGTGAACGTGCAAAACAGGCTCTA
GCGTTCGAACGCACTGATTTCGACCAGGTTCGTTCACTCATGGAAAATAGCGATCGC
TGCCAGGATATACGTAATCTGGCATTTCTGGGGATTGCTTATAACACCCTGTTACGT
ATAGCCGAAATTGCCAGGATCAGGGTTAAAGATATCTCACGTACTGACGGTGGGAG
AATGTTAATCCATATTGGCAGAACGAAAACGCTGGTTAGCACCGCAGGTGTAGAGA
AGGCACTTAGCCTGGGGGTAACTAAACTGGTCGAGCGATGGATTTCCGTCTCTGGTG
TAGCTGATGATCCGAATAACTACCTGTTTTGCCGGGTCAGAAAAAATGGTGTTGCCG
CGCCATCTGCCACCAGCCAGCTATCAACTCGCGCCCTGGAAGGGATTTTTGAAGCAA

CTCATCGATTGATTTACGGCGCTAAGGATGACTCTGGTCAGAGATACCTGGCCTGGT
CTGGACACAGTGCCCGTGTCGGAGCCGCGCGAGATATGGCCCGCGCTGGAGTTTCA
ATACCGGAGATCATGCAAGCTGGTGGCTGGACCAATGTAAATATTGTCATGAACTAT
ATCCGTAACCTGGATAGTGAAACAGGGGCAATGGTGCGCCTGCTGGAAGATGGCGA
TCTCGAGTCTGGAAGTGGTGGTGGAGAACTTGATGAATTGGTATACTTACTAGATGG
GCCAGGTTATGACCCTATACATTGCGATGTAGTGACAAGGGGCGGCAGCCACCTTTT
CAATTTT
